# Comparison of Transformations for Single-Cell RNA-Seq Data

**DOI:** 10.1101/2021.06.24.449781

**Authors:** Constantin Ahlmann-Eltze, Wolfgang Huber

## Abstract

The count table, a numeric matrix of genes × cells, is the basic input data structure in the analysis of single-cell RNA-seq data. A common preprocessing step is to adjust the counts for variable sampling efficiency and to transform them so that the variance is similar across the dynamic range. These steps are intended to make subsequent application of generic statistical methods more palatable. Here, we describe four transformation approaches based on the delta method, model residuals, inferred latent expression state, and factor analysis. We compare their strengths and weaknesses and find that the latter three have appealing theoretical properties. However, in benchmarks using simulated and real-world data, it turns out that a rather simple approach, namely, the logarithm with a pseudo-count followed by principal component analysis, performs as well or better than the more sophisticated alternatives.

**Software:** The R package *transformGamPoi* implementing the delta method- and residuals-based variance-stabilizing transformations is available via Bioconductor. We provide an interactive website to explore the benchmark results at shiny-portal.embl.de/shinyapps/app/08_single-cell_transformation_benchmark.

---

Single-cell RNA sequencing count tables are heteroskedastic. In particular, counts for highly expressed genes vary more than for lowly expressed genes. Accordingly, a change in a gene’s counts from 0 to 100 between different cells is more relevant than, say, a change from 1,000 to 1,100. Analyzing heteroskedastic data is challenging because standard statistical methods typically perform best for data with uniform variance.

One approach to handle such heteroskedasticity is to explicitly model the sampling distributions. For data derived from unique molecular identifiers (UMIs), a theoretically and empirically well-supported model is the Gamma-Poisson distribution^1^ (Grün et al., 2014; Svensson, 2020; Kharchenko, 2021), but parameter inference can be fiddly and computationally expensive (Townes, 2019; Ahlmann-Eltze and Huber, 2020). An alternative choice is to use variance-stabilizing transformations as a preprocessing step and subsequently use the many existing statistical methods that implicitly or explicitly assume uniform variance for best performance (Amezquita et al., 2020; Kharchenko, 2021).

Variance-stabilizing transformations based on the delta method (Dorfman, 1938) promise an easy fix for heteroskedasticity if the variance predominantly depends on the mean. Instead of working with the raw counts *Y*, we apply a non-linear function *g*(*Y*) designed to make the variances (and possibly, higher moments) more similar across the dynamic range of the data (Bartlett, 1947). The Gamma-Poisson distribution with mean *μ* and overdispersion *α* implies a quadratic mean-variance relation 𝕍ar[*Y*] = *μ* + *αμ*^2^. Here, the Poisson distribution is the special case with *α* = 0, and *α* can be considered a measure of additional variation on top of the Poisson. Given such a mean-variance relation, applying the delta method produces the variance-stabilizing transformation

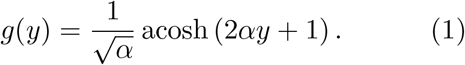

See Appendix B.1 for the derivation. Practitioners often use a more familiar functional form, the shifted logarithm

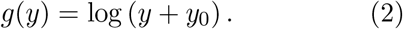

This approximates Eq. (1), in particular if the pseudo-count is *y*_0_ = 1*/*(4*α*) (Appendix B.2).

An additional requirement is posed by experimental variations in sampling efficiency and different cell sizes (Lun et al., 2016), which manifest themselves in varying total numbers of UMIs per cell. Commonly, a so-called size factor *s* is determined for each cell, and the counts are divided by it before applying the variance-stabilizing transformation: e.g., log(*y/s* + *y*_0_) (Love et al., 2014; Amezquita et al., 2020; Borella et al., 2022). There is a variety of approaches to estimate size factors from the data. Conventionally, they are scaled to be close to 1, e. g., by dividing them by their mean, such that the range of the adjusted counts is about the same as that of the raw counts. The simplest estimate of the size factor for cell *c* is

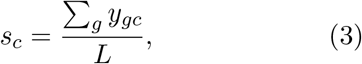

where the numerator is the total number of UMIs for cell *c, g* indexes the genes, and *L* = (#cells)^−1^ ∑_*gc*_ *y*_*gc*_ is the average across all cells of these numerators.

Sometimes, a fixed value is used instead for *L*. For instance, Seurat uses *L* = 10 000, others have used *L* = 10^6^ (Luecken and Theis, 2019), calling the resulting values *y*_*gc*_*/s*_*c*_ *counts per million* (CPM). Even though the choice of *L* may seem arbitrary, it matters greatly. For example, for typical droplet-based single-cell data with sequencing depth of ∑ _*g*_ *y*_*gc*_ ≈ 5 000, using *L* = 10^6^ and then transforming to log(*y*_*gc*_*/s*_*c*_ + 1) is equivalent to setting the pseudo-count to *y*_0_ = 0.005 in Eq. (2). This amounts to assuming an overdispersion of *α* = 50, based on the relation between pseudo-count and overdispersion explained in Appendix B.2. That is two orders of magnitude larger than the overdispersions seen in typical single-cell datasets. In contrast, using the same calculation, Seurat’s *L* = 10 000 implies a pseudo-count of *y*_0_ = 0.5 and an overdispersion of *α* = 0.5, which is closer to overdispersions observed in real data. Yet, choosing *L* or *y*_0_ is unintuitive. Instead, we recommend parameterizing the shifted logarithm transformation in terms of the typical overdispersion, using the relation *y*_0_ = 1*/*(4*α*) motivated above.

**what is the delta method?**

The delta method is a way to find the standard deviation of a transformed random variable.

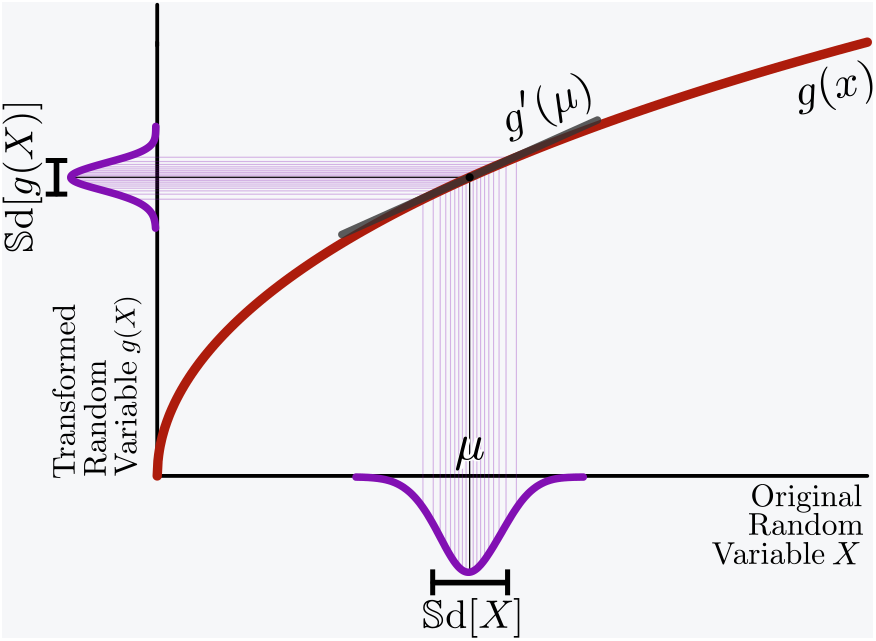

If we apply a differentiable function *g* to a random variable *X* with mean *μ*, the standard deviation of the transformed random variable *g*(*X*) can be approximated by

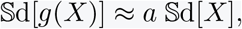

where *a* = |*g*′ (*μ*)| is the slope of *g* at *μ*.

Now consider a set of random variables *X*_1_, *X*_2_, … whose variances and means are related through some function *v*, i. e., 𝕍ar[*X*_*i*_] = *v*(*μ*_i_), or equivalently 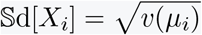. Then we can find a variance-stabilizing transformation *g* by requiring constant standard deviation, 𝕊d[*g*(*X*_*i*_)] = const., which using the above approximation becomes

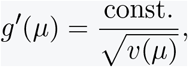

and can be solved by integration.

Hafemeister and Satija (2019) suggested a different approach to variance stabilization based on Pearson residuals

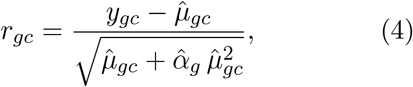

where 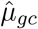 and 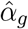 come from fitting a Gamma-Poisson generalized linear model,

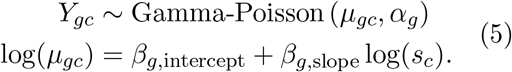

Here, *s*_*c*_ is again the size factor for cell *c*, and *β*_*g*,intercept_ and *β*_*g*,slope_ are intercept and slope parameters for gene *g*. Note that the denominator in Eq. (4) is the standard deviation of a Gamma-Poisson random variable with parameters 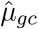 and 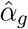.

A third set of transformations infers the parameters of a postulated generative model, aiming to estimate so-called latent gene expression values based on the observed counts. A prominent instance of this approach is *Sanity*, a fully Bayesian model for gene expression (Breda et al., 2021). It infers latent gene expression using a method that resembles a variational mean-field approximation for a log-normal Poisson mixture model. Sanity comes in two flavors: *Sanity Distance* calculates the mean and standard deviation of the posterior distribution of the logarithmic gene expression; based on these, it calculates all cell-by-cell distances, from which it can find the *k* nearest neighbors of each cell. *Sanity MAP* (short for *maximum a posteriori*) ignores the inferred uncertainty and returns the maximum of the posterior as the transformed value. A related tool is *Dino*, which fits mixtures of Gamma-Poisson distributions and returns random samples from the posterior (Brown et al., 2021). *Normalisr* is a tool primarily designed for frequentist hypothesis testing (Wang, 2021), but as it infers logarithmic latent gene expression, it might also serve as a generic preprocessing method. *Normalisr* returns the minimum mean square error estimate for each count assuming a binomial generative model.

In this work, we analyze transformations for preprocessing UMI-based single-cell RNA-seq data based on each of these approaches. We will first contrast the conceptual differences between them. In a second part, we benchmark the empirical performance of all approaches and provide guidelines for practitioners to choose among the methods. In the benchmarks, we also include a fourth preprocessing approach that is not transformation-based and directly produces a low-dimensional latent space representation of the cells: factor analysis for count data based on the (Gamma−)Poisson sampling distribution. An early instance of this approach, called *GLM PCA*, was presented by Townes (2019) and applied to biological data by Townes et al. (2019). Recently, Agostinis et al. (2022) presented an optimized implementation called *NewWave*.

## Results

There are multiple *flavors* for each of the four approaches:

- Among the delta method-based variance-stabilizing transformations, we considered the acosh transformation Eq. (1), the shifted logarithm Eq. (2) with pseudo-count *y*_0_ = 1 or *y*_0_ = 1*/*(4*α*), and the shifted logarithm with counts per million (CPM). In addition, we tested the shifted log transformation with highly variable gene selection (HVG), z-scoring (Z), and rescaling the output as suggested by Booeshaghi et al. (2022).
- Among the residuals-based variance-stabilizing transformations, we considered the clipped and unclipped Pearson residuals (implemented by *sctransform* and *transformGamPoi*) and randomized quantile residuals. In addition, we tested the clipped Pearson residuals with highly variable gene selection, z-scoring, and an analytical approximation to the Pearson residuals suggested by Lause et al. (2021).
- Among the latent gene expression-based transformations (abbreviated Lat. Expr.), we considered *Sanity Distance* and *Sanity MAP, Dino*, and *Normalisr*.
- Among the count-based factor analysis models (abbreviated Count), we considered *GLM PCA* and *NewWave*.

Lastly, we include two methods as negative controls in our benchmarks (abbreviated Neg.), for which we expect poor performance: the raw untransformed counts (*y*) and the raw counts scaled by the size factor (*y/s*).

### Conceptual differences

A known problem for variance-stabilizing transformations based on the delta method derives from the size factors. Fig. 1A shows the first two principal components of a homogeneous solution of droplets encapsulating aliquots from the same RNA (Svensson et al., 2017) for representative instances of the delta method-, residuals- and latent expression-based transformation approaches. (Suppl. Fig. S1 shows the results for all transformations.) Despite the size factor scaling, after the delta method-based transformation, the size factor remained a strong variance component in the data. In contrast, the other transformations better mixed droplets with different size factors. Intuitively, the trouble for the delta method-based transformation stems from the fact that the division of the raw counts by the size factors scales large counts from droplets with large size factors and small counts from droplets with small size factors to the same value. This violates the assumption of a common mean-variance relationship. In Appendix B.3, we dissect this phenomenon more formally.

**Figure 1.**
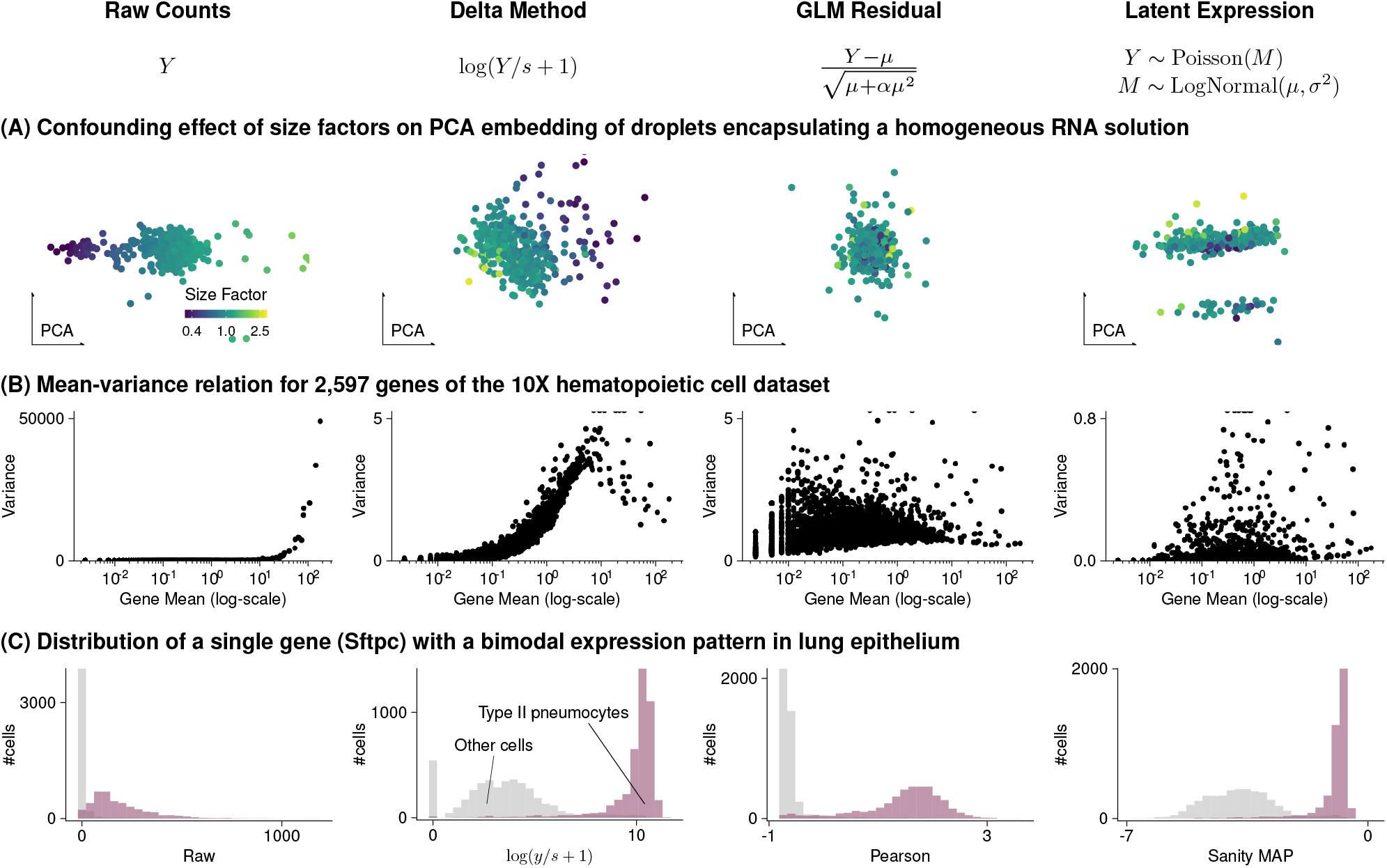
Conceptual differences between variance-stabilizing transformations. The four columns of this figure correspond to raw counts and transformation by shifted logarithm, clipped Pearson residuals, and Sanity MAP. (A) Scatter plot of the first two principal components of data from droplets encapsulating a homogeneous RNA solution. Each point corresponds to a droplet and is colored by its size factor. (B) Scatter plot of the mean-variance relation, where each point is a gene from a human hematopoietic cell dataset. Note that the y-axis range differs between transformations, and outliers are plotted on the edge of the plot. (C) Histogram of the transformed values for Sftpc, a marker for type II pneumocytes, that has a bimodal gene expression in mouse lung epithelium. Details on the data are in Suppl. Table S1.

One of the motivations stated by Hafemeister and Satija (2019) for the Pearson residuals-based variance-stabilizing transformation is that the delta method-based transformations fail to stabilize the variance of lowly expressed genes. Warton (2018) provided a theoretical explanation for this fact. Indeed, Fig. 1B shows that the variance after transformation with a delta method-based variance-stabilizing transformation was practically zero for genes with a mean expression of less than 0.1. In contrast, after residuals-based transformation, the variance showed a weaker dependence on mean expression, save very lowly expressed genes whose variance is limited by the clipping step (compare Pearson and Pearson (no clip) in Suppl. Fig. S2). The results of the latent expression-based transformations were diverse, reflecting that these methods are not directly concerned with stabilizing the variance. Individual patterns ranged from higher variance for lowly expressed genes (*Sanity Distance* and *Normalisr*) to the opposite trend for *Dino* (Suppl. Fig. S2).

A peculiarity of the Pearson residuals is their behavior if a gene’s expression strongly differs between cell subpopulations. Fig. 1C shows a bimodal expression pattern of Sftpc, a marker for type II pneumocytes. Unlike the transformations based on the delta method or latent expression models, the Pearson residuals are an affine-linear transformation per gene (Eq. (4)) and thus cannot shrink the variance of the high-expression subpopulation more than that of the low-expression subpopulation (compare the Pearson residuals with *y/s* in Suppl. Fig. S3). This can affect visualizations of such genes and, in principle, other analysis tasks such as detection of marker genes or clustering and classification of cells.

An alternative is to combine the idea of delta method-based variance-stabilizing transformations with the generalized linear model residuals approach by using non-linear residuals. We considered randomized quantile residuals (Dunn and Smyth, 1996). (Suppl. Fig. S4 shows how they are constructed.) Like Pearson residuals, randomized quantile residuals stabilized the variance for small counts (Suppl. Fig. S2), but in addition, they also stabilized the within-group variance if a gene’s expression strongly differed across cells (Suppl. Fig. S3).

Such conceptual differences of the transformation approaches are important to understand when applying them to novel data types or when developing new transformations; but for most practitioners, empirical performance will be of primary interest. We look at this in the next section.

### Benchmarks

There is no context-free measure of success for a preprocessing method, as it is contingent on the objectives of the subsequent analysis. For instance, if interest lies in identification of cell type-specific marker genes, one could assess the shape of distributions, such as in Fig. 1C, or the performance of a supervised classification method. Here, we considered the objective that arguably has been the main driver of single-cell RNA-Seq development and applications so far: understanding the variety of cell types and states in terms of a lower-dimensional mathematical structure, such as a planar embedding, a clustering, trajectories, branches, or combinations thereof. For all of these, one can consider the *k*-nearest neighbor (*k*-NN) graph as a fundamental data structure that encodes essential information. The next challenge is then the definition of “ground truth”. We designed our benchmarks upon reviewing previous benchmarking approaches. For instance, Breda et al. (2021) and Lause et al. (2021) employed synthetic or semi-synthetic data. This is operationally attractive, but it is difficult to be certain about biological relevance. Hafemeister and Satija (2019) and Lause et al. (2021) used qualitative inspection of non-linear dimension reduction plots. This can be informative, but is difficult to scale up and make objective. Germain et al. (2020) compared how well the transformations recovered *a priori* assigned populations, defined either through FACS or by mixing different cell lines. This is conceptually clean, but restricts analysis to a limited range of data sets that also may only offer a caricature view of cell diversity.

For all our benchmarks, we applied the transformations to the raw counts of each dataset listed below, computed a lower-dimensional representation of the cells using principal component analysis (PCA), identified the *k* nearest neighbors of each cell as measured by Euclidean distance, and, finally, computed the overlap of the thus obtained *k*-NN graph with a reference *k*-NN graph (see Methods for details). We performed these three benchmarks:

#### Consistency

We downloaded ten 10X datasets from the GEO database. Since there was no formal ground truth, we measured the consistency of the results (a necessary, although not sufficient, condition for their goodness) by splitting the genes of each dataset into two disjoint subsets.

#### Simulation

We used four different previously published simulation frameworks and one adapted by us to generate a diverse collection of datasets for which we had full access to the true *k*-NN graph.

#### Downsampling

We used five deeply sequenced datasets based on mcSCRB and Smart-seq3 (details in Suppl. Table S2), which we down-sampled to sequencing depths typical for the 10X technology. We postulated that a proxy for ground truth could be constructed from the *k*-NN graph inferred from the deeply sequenced data intersected across all transformations which we call reliable nearest neighbors. To our knowledge, this work presents the first instance of such an approach.

Suppl. Table S3 and Suppl. Figs. S5 and S6 give an overview over the datasets.

We tested 22 transformations—where applicable with an overdispersion fixed to 0, 0.05, and a gene-specific estimate from the data—across four to eight settings for the number of dimensions of the PCA and measured the overlap with *k* = 10, 50, and 100 nearest neighbors. In total, we collected more than 61 000 data points. In addition to the results highlighted in the following, we provide an interactive website with all results for all tested parameter combinations.

Fig. 2 shows the aggregated results for the three benchmarks for *k* = 50. Similar results were obtained for *k* = 10 and *k* = 100, shown in Suppl. Fig. S7.

**Figure 2.**
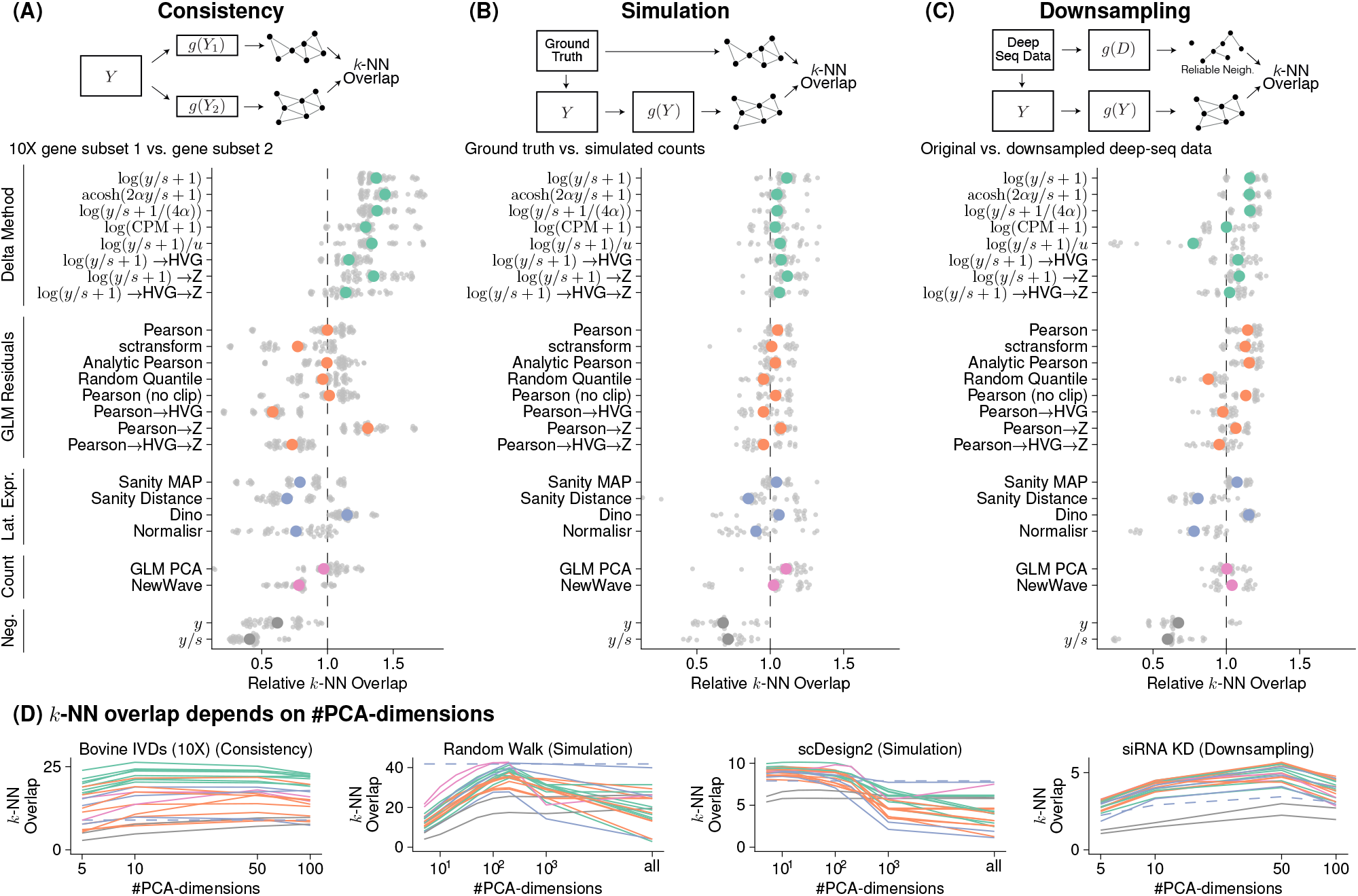
Benchmark results. (A) Overlap between the *k*-nearest neighbors (*k*-NN) inferred separately on two halves of the data. The colored points show the averages across 10 datasets, each with 5 replicate random data splits (small, grey points). (B) Overlap between *k*-NN inferred from simulated data and ground truth, using 5 simulation frameworks and 5 replicates per framework. (C) Overlap between a reference *k*-NN graph (inferred using all transformations on deeply sequenced data and taking the intersection) and the *k*-NN inferred on data downsampled to match typical 10X data (i.e., 5,000 counts per cell) for 5 datasets with 5 replicates each. To compare and aggregate results across the different datasets, Panels A-C show relative overlap, which was computed by dividing, for each dataset, the overlap by its average across all transformations, fixing *k* = 50 and using a dataset-specific number of PCA dimensions (Suppl. Fig. S8 shows the underlying, unaggregated data). (D) Overlap (*y*-axis) as a function of PCA dimensions (*x*-axis); the different transformation types are indicated by the colors, using the same palette as in Panels A-C. The performance of *Sanity Distance* is shown as a dashed line.

In the consistency benchmark, the delta method-based transformations performed better than the other transformations (Fig. 2A).

On the simulated data, the differences between the transformations looked less pronounced in Fig. 2B than for the other two benchmarks. However, this is a result of the aggregated view. For each particular simulation framework, large differences between the transformations appeared, but the results varied from simulation to simulation framework (Suppl. Fig. S8B) and averaged out in the aggregated view.

The results of the downsampling benchmark (Fig. 2C) agreed well with the trends observed in the simulation and the consistency benchmark. This benchmark is particularly informative because the data had realistic latent structures, which were reliably detectable through the high sequencing depth. The downsampling produced data that resembles the more common 10X data in many characteristics: e.g., UMIs per cell, proportion of zeros in the data (Suppl. Tab. S3), and mean-variance relation (Suppl. Fig. S5). The main difference was that the suitable (high sequencing depth per cell) datasets we could access mostly comprised only a few hundred cells, except for the 4 298 cells siRNA KD dataset (Suppl. Fig. S6).

The results in Fig. 2 are on a relative scale, which hides the magnitude of the differences. In Suppl. Fig. S8, we show the underlying results for each dataset on an absolute scale. The range of *k*-NN overlaps was dataset dependent, ranging from 34/50 for the best performing transformation versus 9/50 for the negative control for the SUM149PT cell line consistency benchmark, to 2.9/50 vs. 1.5/50 for the HEK downsampling benchmark. For the latter, the overall small overlaps were due to small sets of reliable nearest neighbors (Suppl. Fig. S9). We also ran a version of the downsampling benchmark that only used the top two transformations per approach (Suppl. Fig. S10), which increased the number of reliable nearest neighbors, and confirmed the trends we saw in the full version.

In addition to the *k*-NN overlap with the ground truth, we also calculated the adjusted Rand index (ARI) and the adjusted mutual information (AMI) for the five simulation frameworks. Suppl. Figs. S11A, B show the aggregated results, which were similar to the results for the *k*-NN overlap (Fig. 2B). Suppl. Figs. S11C, D show that the ARI and AMI had a larger dynamic range than the *k*-NN overlap for datasets with a small number of distinct clusters; however, for datasets with a complex latent structure, the *k*-NN overlap was more informative, which may reflect limitations of ARI and AMI to assess the recovery of gradual changes typical for many biological tissues.

The Random Walk simulation reproduced the benchmark based on which Breda et al. (2021) argued that *Sanity* was the best method for identifying the *k* nearest neighbors of a cell (Fig. 5A of their paper). We found that the delta method-based and residuals-based variance-stabilizing transformations performed as well in this benchmark if we projected the cells to a lower-dimensional representation before constructing the *k*-NN graph. In fact, Fig. 2D shows for four example datasets that the number of dimensions for the PCA was an important determinant of performance. This is because the dimension reduction acts as a smoothener, whose smoothing effect needs to be strong enough to average out uncorrelated noise (i.e., small enough target space dimension), but flexible enough to maintain interesting variation (i.e., large enough target space dimension).

The latent expression-based transformations (except *Normalisr*) and the count-based factor analysis models were computationally more expensive than the delta method- and residuals-based transformations. Fig. 3A shows the CPU times for calculating the transformation and finding the *k* nearest neighbors on the *10X human T helper cell* dataset with 10 064 cells. *Sanity Distance* took particularly long because its distance calculation, which takes into account the uncertainty for the nearest neighbor search, scaled quadratically with the number of cells (Fig. 3B). Across all benchmarks, the computations took 24 years of CPU time, of which the latent expression-based transformations accounted for over 90%. The delta method-based transformations were the fastest, especially if the overdispersion was not estimated from the data. The residuals-based transformations took somewhat more time, except for the analytic approximation of the Pearson residuals, which could be calculated almost as fast as the shifted logarithm. In terms of memory consumption, the delta method-based transformations were most attractive because they retained the sparsity of the data.

**Figure 3.**
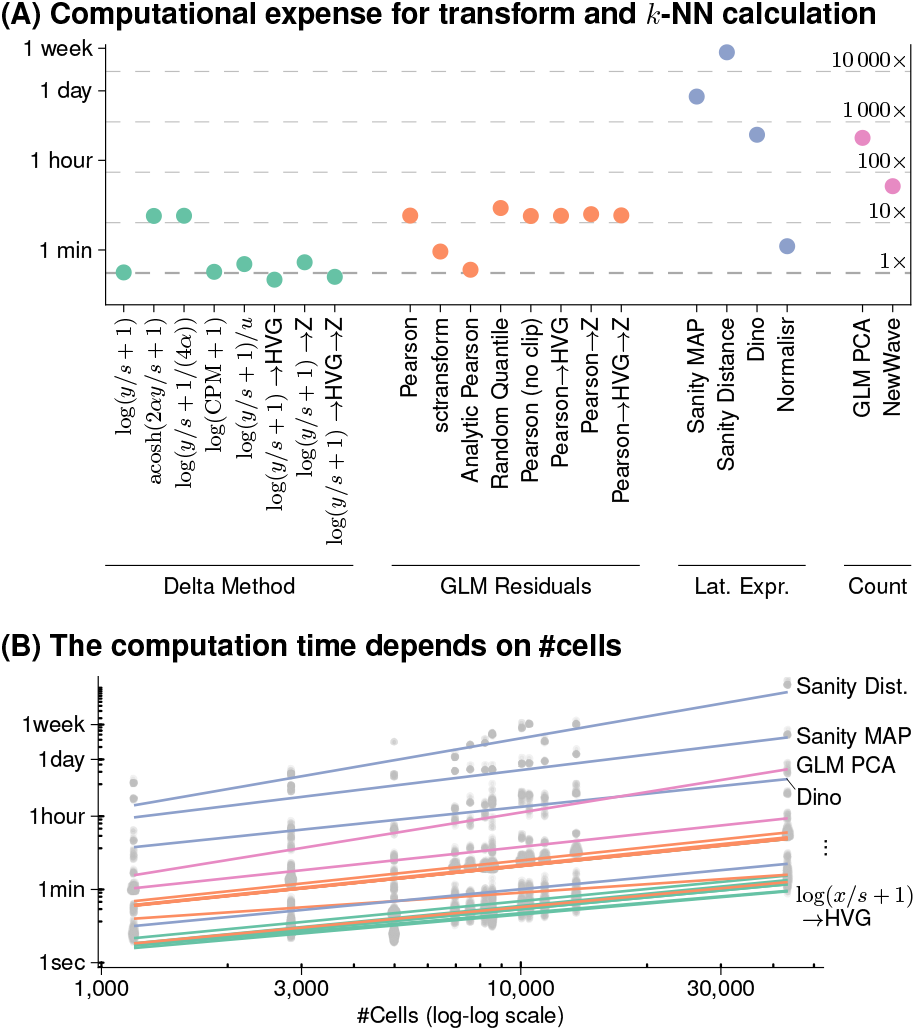
Computational Expense. (A) CPU time needed to calculate the transformation and identify the *k*-NNs for the *10X human T helper cell* dataset. The secondary axis shows the duration relative to the shifted logarithm. (B) Dependence of run time on the number of cells, across datasets, shown on a double-logarithmic scale, with a linear fit. Most transformations have a slope of approximately 1 (i.e., scale linearly), whereas Sanity Distance and GLM PCA have a slope *>* 1.5 which indicates quadratic scaling.

In terms of uncovering the latent structure of the datasets, none of the other transformations consistently outperformed the shifted logarithm (Fig. 4A), one of the simplest and oldest approaches. In fact, when followed by PCA dimension reduction to a suitable target dimension, the shifted logarithm performed better than the more complex latent expression-based transformations across all three benchmarks.

**Figure 4.**
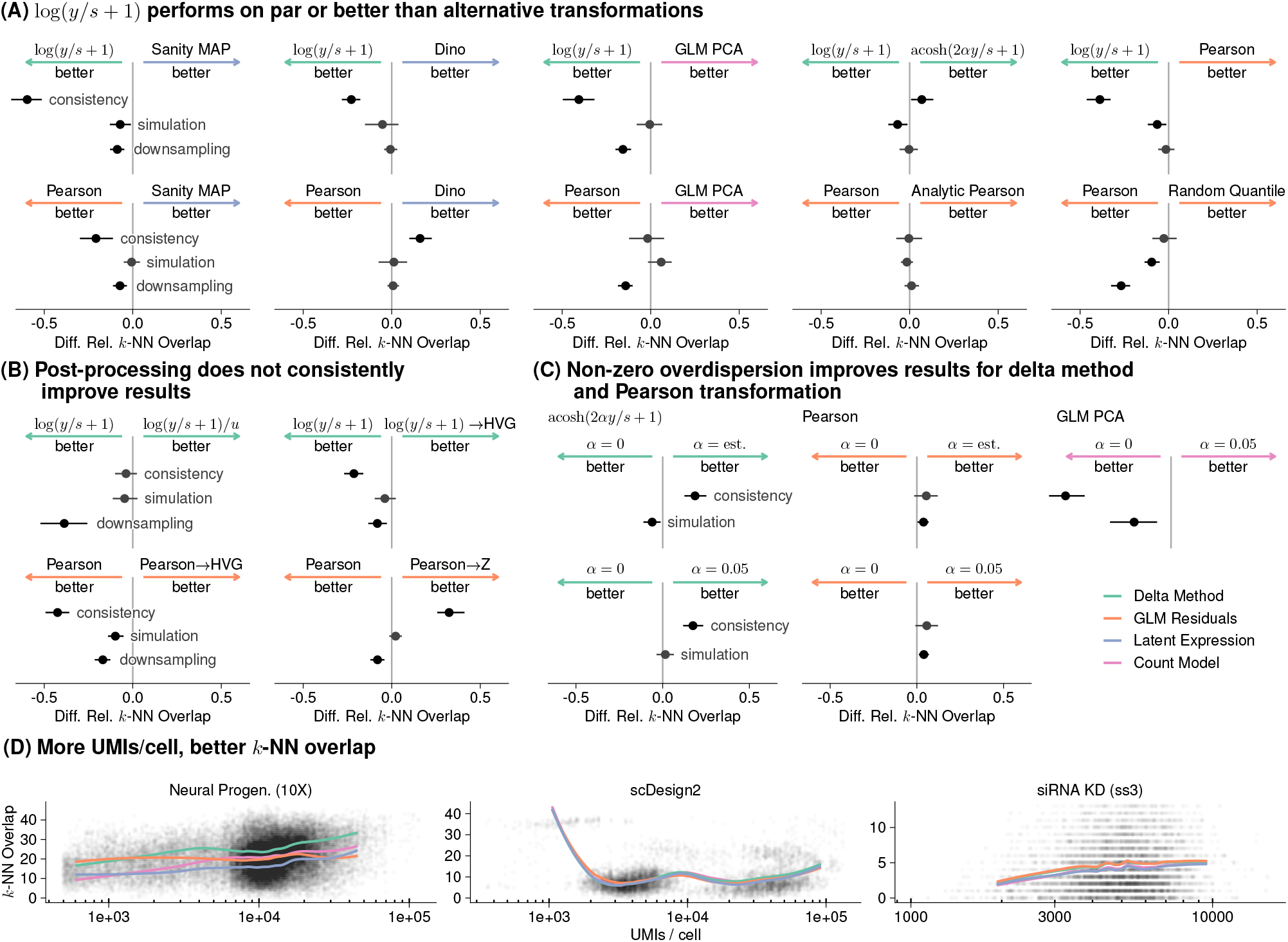
Comparison of selected transformations. (A-C) 95% confidence intervals of the differences of the relative *k*-NN overlap between selected transformations as shown in Fig. 2. (A) Shifted logarithm and Pearson residuals against a selection of the best performing transformations from the other preprocessing approaches. (B) Effect of applying various post-processing methods after applying the shifted logarithm or Pearson residuals transformation. (C) Effect of fixing the overdispersion to 0 or 0.05, or estimating a gene-specific overdispersion from the data. (D) Smoothed line plots of the *k*-NN overlap (y-axis) as a function of the UMIs per cell (x-axis) for the shifted logarithm transformation, the Pearson residuals, Sanity MAP, and GLM PCA colored by the respective transformation approach.

We found no evidence that additional post-processing steps (rescaling the output of the shifted logarithm, selecting highly variable genes, or equalizing the variance of all genes using z-scoring) improved the results for identifying nearest neighbors (Fig. 4B). Lause et al. (2021) and Choudhary and Satija (2022) debated on how to best choose the overdispersion parameter. We found empirically that, for Pearson residuals and the acosh transformation, it is beneficial to have *α >* 0, but saw no clear benefits from estimating this parameter from the input data versus using a generic, fixed value of, say, 0.05 (Fig. 4C).

Lastly, we found that with increasing sequencing depth per cell, all methods generally had a better *k*-NN overlap with the ground truth (Fig. 4D). This makes intuitive sense: with higher sequencing depth, the relative size of the sampling noise is reduced. Based on Fig. 1A, we might assume that delta method-based transformations would perform particularly poorly at identifying the neighbors of cells with extreme sequencing depths; yet on three datasets, the shifted logarithm performed as well for cells with particularly large or small size factors, as for other cells (Fig. 4D). We also considered the performance of the transformations as a function of cluster size (Suppl. Figs. S12-S14); while we see some interesting variation, we do not find that a single transformation performed consistently better or worse for small clusters.

## Discussion

We compared 22 transformations, conceptually grouped into four basic approaches, for their ability to recover latent structure among the cells. We found that one of the simplest approaches, the shifted logarithm transformation log(*y/s*+*y*_0_) with *y*_0_ = 1 followed by PCA, performed surprisingly well. We presented theoretical arguments for using the related acosh transformation or an adaptive pseudo-count *y*_0_ = 1*/*(4*α*), but our benchmarks showed limited performance benefits for these.

We recommend against using counts per million as input for the shifted logarithm. We pointed out that for typical datasets, this amounts to assuming an unrealistically large overdispersion, and in our benchmarks this approach performed poorly compared to applying the shifted logarithm to size factor-scaled counts. We also advise against scaling the results of the shifted logarithm by the sum of the transformed values per cell as, e.g., suggested by Booeshaghi et al. (2022). In our hands (Suppl. Fig. S1), this additional operation failed to remove the confounding effect of the sequencing depth (the authors’ stated motivation for it) and did not improve the *k*-NN recall performance.

The Pearson residuals-based transformation has attractive theoretical properties and, in our benchmarks, performed similarly well as the shifted logarithm transformation. It stabilizes the variance across all genes and is less sensitive to variations of the size factor. The analytic approximation suggested by Lause et al. (2021) is appealing because it worked as well as the exact Pearson residuals but could be calculated faster. However, as seen in Eq. (4), the Pearson residuals-based transformation is affine linear when considered as a function per gene, and this may be unsatisfactory for genes with a large dynamic range across cells. As an alternative, we considered randomized quantile residuals as a non-linear transformation, but found no performance improvement. This result exemplifies that choosing a transformation for conceptual reasons does not necessarily translate into better downstream analysis results.

The use of the inferred latent expression state as a transformation and count-based latent factor models are appealing because of their biological interpretability and mathematical common sense. In particular, Sanity Distance is appealing because it does not have any tunable parameters. However, all these transformations performed worse than the shifted logarithm with a reasonable range of PCA dimensions in our benchmarks and some of the transformations were exceptionally computationally expensive (e.g., the median CPU time of Sanity Distance was 4 500× longer than for the shifted logarithm).

Our results partially agree and disagree with previous studies. Germain et al. (2020) benchmarked many steps of a typical single-cell RNA-seq analysis pipeline, including a comparison of clustering results obtained after different transformations against *a priori* assigned populations. In line with our findings, they reported that dimension reduction was of great importance. They went on to recommended *sctransform* (i.e., Pearson residuals) based on its good performance on the *Zhengmix4eq* dataset, which is a mixture of PBMCs sorted by surface markers using flow cytometry. However, it is not clear how generalizable this result is, and our benchmarks do not support such a singling out of that method. Lause et al. (2021) considered the related *Zhengmix8eq* dataset, into which they implanted a synthetic rare cell type by copying 50 B-cells and appending 10 genes exclusively expressed in the synthetic cell type. They used *k*-NN classification accuracy of the cell type averaged per cell type (macro F1 score, Fig. 5c of their paper) and averaged over all cells (online version of Fig. 5c). They found a performance benefit for the Pearson residuals over the shifted logarithm with the macro F1 score, but similar performance with regard to overall accuracy. The macro F1 score emphasizes the performance difference for the synthetic cell type, which appears somewhat construed and might not be a good model for most biologically relevant cell type and state differences. Instead of comparing clustering results to discrete cell type assignments, we have focused on the inference of the *k* nearest neighbors of each cell, with the expectation that this enables consideration of more subtle latent structures than well-separated, discrete cell types.

Pearson residuals- and delta method-based transformations weight genes differently; e.g., Pearson residuals put more weight on lowly expressed genes than the delta method (Fig. 1B). This can lead to different downstream results, but our benchmarks did not indicate that any particular weighting is generally better; only that the delta method-based transformation produced more consistent results on the 10X datasets.

We did not evaluate the impact of alternative size factor estimators. We also did not consider how suitable a transformation is for marker gene selection, because we are not aware of a suitable metric to determine success, as the utility of a marker gene hinges on its biological interpretability. For a recent effort to compare different marker gene selection methods, see Pullin and McCarthy (2022).

Considerable effort has been invested in the space of preprocessing methods for single-cell RNA-seq data. Somewhat to our surprise, the shifted logarithm still performs among the best for preprocessing, but crucially only if combined with a dimensionality reduction method like PCA and an appropriate number of latent dimensions.

## Availability

An R package that implements the delta method- and residuals-based variance-stabilizing transformations is available on bioconductor.org/packages/transformGamPoi/. The code to generate the figures is available on github.com/const-ae/transformGamPoiPaper. We provide an interactive website to explore the benchmark results at shiny-portal.embl.de/shinyapps/app/08_single-cell_transformation_benchmark. All datasets used in this manuscript are listed in Suppl. Table S1 and S2.

## Acknowledgments

We thank Dr. Simon Anders for extensive discussions about variance-stabilizing transformations and how to benchmark preprocessing methods. We thank the three anonymous reviewers, Prof. Dr. Erik van Nimwegen, and Dr. Dmitry Kobak, whose feedback on an earlier version helped to improve the manuscript.

## Methods

### Transformations

We compared 22 transformations that can be grouped into four approaches.

The delta method-based transformations were: the shifted logarithm (log(*y/s* + 1)), the acosh transformation (acosh(2*αy/s* + 1)), the shifted logarithm with pseudo-count dependent on the overdispersion (log(*y/s* + 1*/*(4*α*))), the shifted logarithm with counts-per-million (log(CPM + 1)), the shifted logarithm with subsequent size normalization as suggested by Booeshaghi et al. (2022) (*x*_*gc*_*/u*_*c*_, where *x*_*gc*_ = log(*y*_*gc*_*/s*_*c*_ + 1) and *u*_*c*_ = ∑_*g*_ *x*_*gc*_), the shifted logarithm with subsequent highly variable gene selection (log(*y/s* + 1) → HVG), the shifted logarithm with subsequent z-scoring per gene (log(*y/s* + 1) → Z), the shifted logarithm with subsequent highly variable gene selection and z-scoring per gene (log(*y/s* + 1) → HVG → Z). For all composite transformations, we first calculated the variance stabilizing transformation, then chose the highly variable genes and used the results without recalculating the variance stabilizing transformation.

To retain the sparsity of the output also if the pseudo-count *y*_0_ ≠ 1, *transformGamPoi* uses the relation

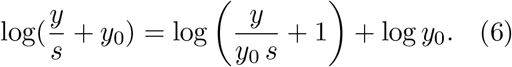

Subtracting the constant log *y*_0_ from this expression does not affect its variance stabilizing properties, but has the desirable effect that data points with *y* = 0 are mapped to 0.

The residuals-based transformations were: Pearson residuals implemented with the *transformGamPoi* package where each residual is clipped to be within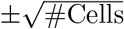, as suggested by Hafemeister and Satija (2019) (Pearson), Pearson residuals with clipping and additional heuristics implemented by *sctransform* Version 2 (sctransform), an analytic approximation to the Pearson residuals with clipping suggested by Lause et al. (2021) (Analytic Pearson), randomized quantile residuals implemented by *transformGamPoi* (Random. Quantile), Pearson residuals without clipping implemented by *transformGamPoi* (Pearson (no clip)), Pearson residuals with clipping and subsequent highly variable gene selection (Pearson → HVG), Pearson residuals with clipping and subsequent z-scoring per gene (Pearson → Z), Pearson residuals with clipping and subsequent highly variable gene selection and z-scoring per gene (Pearson → HVG → Z). For each composite Pearson residual transformation (i.e., with HVG. and / or z-scoring), we used the *transformGamPoi* implementation.

The latent expression-based transformations were: *Sanity* with point estimates for the latent expression (*Sanity MAP*) and with calculation of all cell-by-cell distances taking into account uncertainty provided by the posteriors (*Sanity Distance*), *Dino* as provided in the corresponding R package, and *Normalisr* with variance normalization, implemented in Python, which we called from R using the *reticulate* package.

The count-based factor analysis models were: *GLM PCA* using the Poisson model and the Gamma-Poisson model with *α* = 0.05. In the figures, we show the results for the Poisson model unless otherwise indicated. We used the *avagrad* optimizer. We ran *NewWave* with 100 genes for the mini-batch overdispersion estimation.

For the delta method-based transformations and the residuals-based transformations calculated with the *transformGamPoi* package, we calculated the size factor *s* using Eq. (3).

We defined highly variable genes (HVG) as the 1 000 most variable genes based on the variance of the transformed data.

For z-scoring, we took the transformed values *x*_*gc*_ = *g*(*y*_*gc*_) and computed 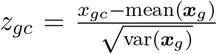, where mean and variance are the empirical mean and variance taken across cells.

In the overview figures (Fig. 2, 3, and 4), we use a gene-specific overdispersion estimate for all residuals-based transformations and for the delta method-based transformations which can handle a custom overdispersion; for *GLM PCA*, we use *α* = 0, because these settings worked best for the respective transformations. The latent expression-based transformations and NewWave do not support custom overdispersion settings.

### Conceptual differences

For the visualization of the residual structure after adjusting for the varying size factors, we chose a control dataset of a homogeneous RNA solution encapsulated in droplets (Svensson et al., 2017). We filtered out RNAs that were all zero and plotted the first two principal components. Where applicable, we used the gene-specific overdispersion estimates. For visualizing the results of Sanity Distance, instead of the PCA, we used multidimensional scaling of the cell-by-cell distance matrix using R’s *cmdscale* function. We calculated the canonical correlation using R’s *cancor* function on the size factors and the first 10 dimensions from PCA and multi-dimensional scaling.

The plots of the mean-variance relation are based on the *10X human hematopoietic cell* dataset (Bulaeva et al., 2020). Where applicable, we used the gene specific overdispersion estimates. The panel of Sanity Distance shows the variance of samples drawn from a normal distribution using the inferred mean and standard deviation.

For the mouse lung dataset (Angelidis et al., 2019), we filtered out cells with extreme size factors (0.1*s*_*median*_ *< s*_*c*_ *<* 10*s*_*median*_, where *s*_*median*_ is the median size factor). We also removed cells that did not pass the *scran* quality control criterion regarding the fraction of reads assigned to mitochondrial genes. To account for the fact that some transformations share information across genes, we applied all transformations to the 100 most highly expressed genes and three genes (Sftpc, Scgb1a1, Ear2) known to be differentially expressed in some cell types according to the assignment from the original publication.

### Benchmark

The benchmarks were executed using a custom work scheduler for slurm written in R on CentOS7 and R 4.1.2 with Bioconductor version 3.14. The set of R packages used in the benchmark with exact version information was stored using the *renv* package and is available from the GitHub repository.

#### *k*-NN identification and dimensionality reduction

To calculate the PCA, we used the *irlba* package. To infer the *k* nearest neighbors, we used *annoy*, which implements an approximate nearest neighbor search algorithm. To calculate the t-SNEs, which we only used for visualization, we used the *Rtsne* package on data normalized with the shifted logarithm with a pseudo-count of 1.

#### Consistency Benchmark

We downloaded ten single-cell datasets listed in the gene expression omnibus database (GEO) browser after searching for the term *mtx* on 2021-10-14. All Datasets are listed in Suppl. Tab. S2. To measure the consistency of the transformations, we randomly assigned each gene to one of two groups and processed the two resulting data subsets separately. We calculated the consistency as the mean overlap of the *k* nearest neighbors for all cells.

#### Simulation Benchmark

We used five frameworks to simulate single-cell counts in R: we ran *dyngen* (Cannoodt et al., 2021) using a consecutive bifurcating mode and the default parameters otherwise. We ran *muscat* (Crowell et al., 2020) with 4 clusters, a default of 30% differentially expressed genes with an average log-fold change of 2, and a decreasing relative fraction of log-fold changes per cluster. We ran *scDesign2* (Sun et al., 2021) with the *10X human hematopoietic cell* dataset as the reference input with a copula model and a Gamma-Poisson marginal distribution. We simulated the *Random Walk* by translating the Matlab code of Breda et al. (2021) to R and using the data by Baron et al. (2016) as a reference. For the *Linear Walk*, we adapted the Random Walk simulation and, instead of following a random walk for each branch, we interpolated the cells linearly between a random start and end point. For both benchmarks, we used a small non-zero overdispersion of *α* = 0.01 to mimic real data.

With each simulation framework, we knew which cells were, in fact, the *k* nearest neighbors to each other. We calculated the overlap as the mean overlap of this ground truth with the inferred nearest neighbors on the simulated counts for all cells. Furthermore, we calculated the adjusted Rand index (ARI) and adjusted mutual information (AMI) by clustering the ground truth and the transformed values with the graph-based walktrap clustering algorithm from the *igraph* package.

#### Downsampling Benchmark

We searched the literature for single-cell datasets with high sequencing depth and found five (one from mcSCRB, four from Smart-seq3) that had a sequencing depth of more than 50 000 UMIs per cell on average. We defined reliable nearest neighbors as the set of *k* nearest neighbors of a cell that were identified with all 22 transformations on the deeply sequenced data (excluding the two negative controls). We used the *downsampleMatrix* function from the *scuttle* package to reduce the number of counts per cell to approximately 5 000, a typical value for 10X data. We considered only one setting for the overdispersion per transformation (instead of allowing multiple overdispersion settings for some transformations as in the other benchmarks). We ran all transformations, which supported the setting, with a gene-specific overdispersion estimate (except *GLM PCA* which performed better with an overdispersion fixed to 0). Finally, we computed the mean overlap between the *k* nearest neighbors identified on the downsampled data with the set of reliable nearest neighbors for all cells with more than one reliable nearest neighbor.

#### *k*-NN Overlap

For all three benchmarks, we calculated overlaps between pairs of *k* nearest neighbor graphs. Denoting their #cell × #cell adjacency matrices (i.e., a matrix of zeros and ones, where an entry is is one if a cell *d* is among the *k* nearest neighbors of cell *c*) by *N* ^1^ and *N* ^2^, we defined their overlap as

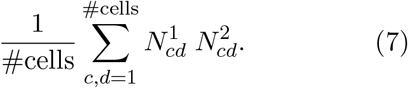

## Supplementary Tables

**Suppl. Table S1:**
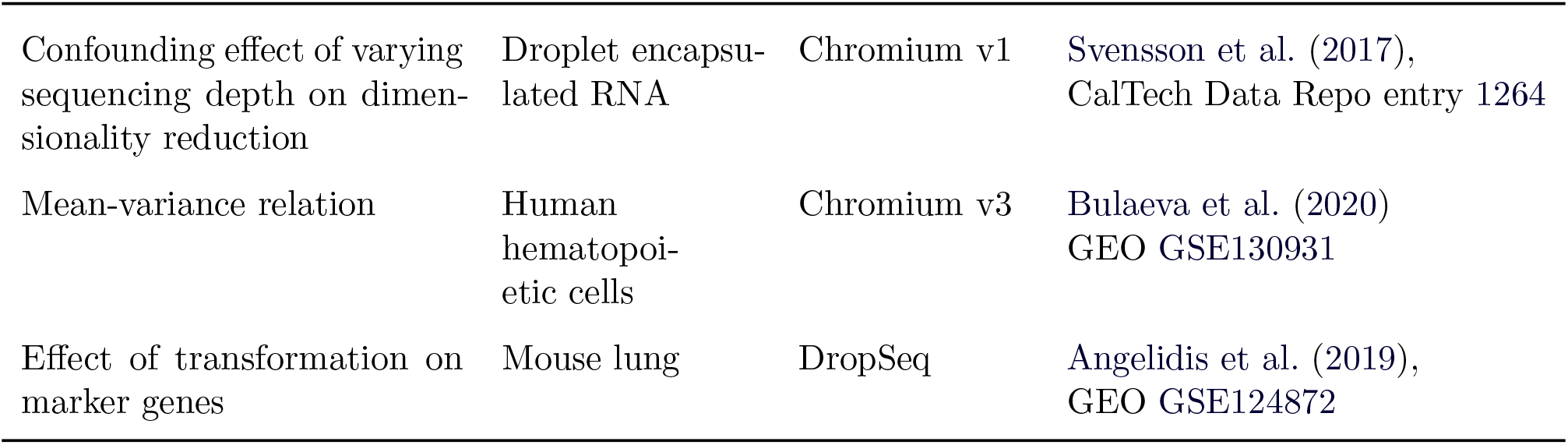
Datasets used to illustrate the conceptual differences between transformations.

**Suppl. Table S2:**
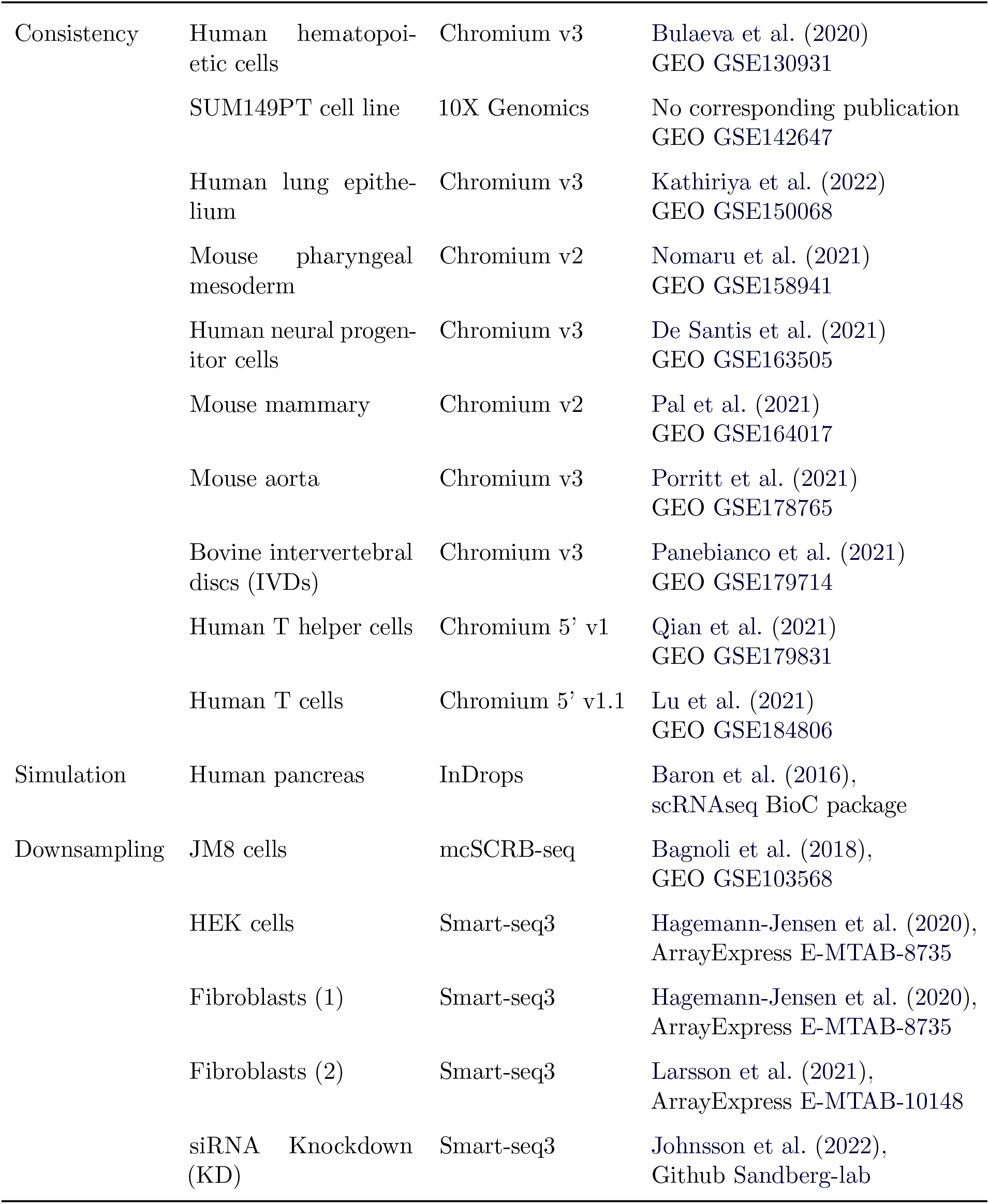
Datasets used to benchmark the performance differences between transformations.

**Suppl. Table S3:** Overview of the datasets used for the benchmark. The *#Genes* and *#Cells* columns show the number of rows and columns in the count matrix after filtering out rows and columns for which all values were zero. *Perc. Zeros* shows what fraction of all values were 0. *99% Quant* shows the 99% quantile of the counts. *Overdisp*. shows the global overdispersion estimate with *glmGamPoi*.

|  | #Cells | #Genes | Perc. Zeros | 99% Quant | UMI/cell | Overdisp. |
| --- | --- | --- | --- | --- | --- | --- |
| **Consistency** |  |  |  |  |  |  |
| Hematopoietic Cells | 2,838 | 21,398 | 87% | 12 | 5,020 | 0.33 |
| SUM149PT Cells | 1,196 | 25,231 | 74% | 35 | 54,900 | 0.14 |
| Lung Epithelium | 11,407 | 20,728 | 90% | 5 | 7,730 | 0.17 |
| Pharyngeal Mesoderm | 7,581 | 19,939 | 79% | 19 | 21,700 | 0.12 |
| Neural Progenitors | 13,572 | 25,711 | 87% | 7 | 11,500 | 0.31 |
| Mouse Mammary | 6,969 | 19,757 | 89% | 6 | 6,970 | 0.24 |
| Mouse Aorta | 10,477 | 20,020 | 86% | 8 | 9,420 | 0.89 |
| Bovine IVDs | 8,231 | 17,464 | 90% | 6 | 3,940 | 1.20 |
| T Helper Cells | 10,064 | 21,153 | 83% | 15 | 19,300 | 0.33 |
| T Cells | 43,283 | 23,978 | 92% | 4 | 5,360 | 0.53 |
| **Simulation** |  |  |  |  |  |  |
| Dyngen | 5,000 | 995 | 75% | 3 | 291 | 0.20 |
| Linear Walk | 8,569 | 17,130 | 90% | 5 | 4,340 | 2.20 |
| muscat | 5,000 | 999 | 63% | 22 | 1,830 | 0.98 |
| Random Walk | 8,569 | 17,192 | 90% | 5 | 4,820 | 2.60 |
| scDesign2 | 2,838 | 16,199 | 82% | 15 | 5,170 | 0.35 |
| **Downsampling (original)** |  |  |  |  |  |  |
| mcSCRB | 249 | 16,864 | 57% | 48 | 59,000 | 0.47 |
| Fibroblasts | 369 | 16,535 | 45% | 224 | 199,000 | 0.82 |
| Fibroblasts 2 | 737 | 18,682 | 48% | 181 | 197,000 | 0.33 |
| HEK | 339 | 18,746 | 63% | 38 | 56,100 | 0.15 |
| siRNA KD | 4,298 | 18,956 | 56% | 106 | 122,000 | 0.36 |
| **Downsampling (reduced)** |  |  |  |  |  |  |
| mcSCRB | 249 | 16,864 | 87% | 5 | 5,020 | 0.32 |
| Fibroblasts | 369 | 16,535 | 85% | 6 | 5,020 | 0.19 |
| Fibroblasts 2 | 737 | 18,682 | 88% | 5 | 5,020 | 0.13 |
| HEK | 339 | 18,746 | 89% | 4 | 5,140 | 0.11 |
| siRNA KD | 4,298 | 18,956 | 88% | 5 | 4,990 | 0.23 |

### A Supplementary Figures

**Suppl. Figure S1:**
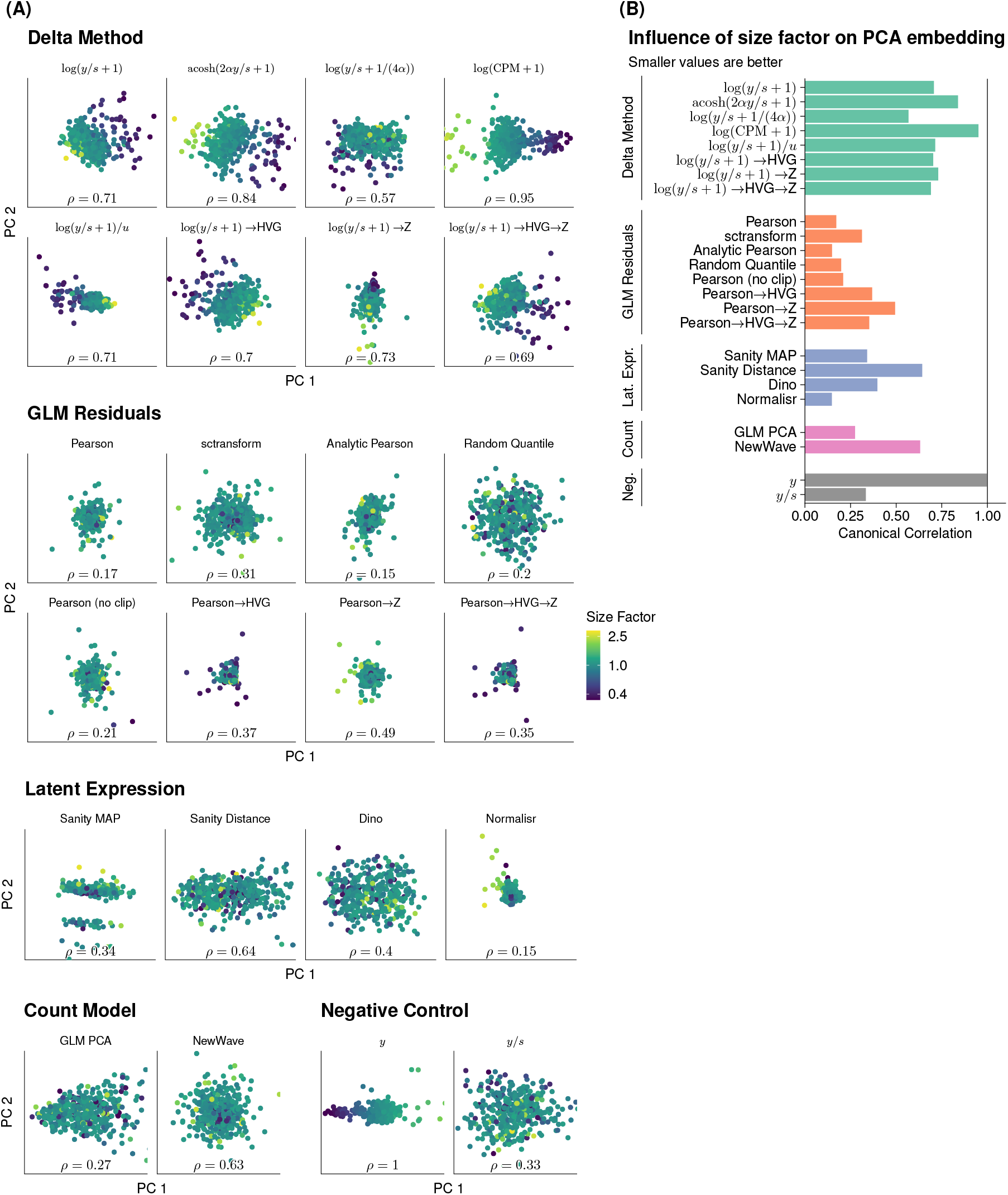
Scatter plot of the first two principal components of the transformed data colored by the sequencing depth (expressed as a normalized size factor on a logarithmic scale) per cell. The data are from droplets that encapsulate a homogeneous RNA solution, and thus the only variation is due to technical factors like sequencing depth (Svensson et al., 2017). The annotation at the bottom of the plot shows the canonical correlation coefficient *ρ* (Hotelling, 1936) between the size factor and the first ten principal components. A lower canonical correlation means that the variance-stabilizing transformation more successfully adjusts for the varying size factors; a canonical correlation of *ρ* = 1 means that the ordering of the cells along some direction in the first 10 PCs is entirely determined by the size factor.

**Suppl. Figure S2:**
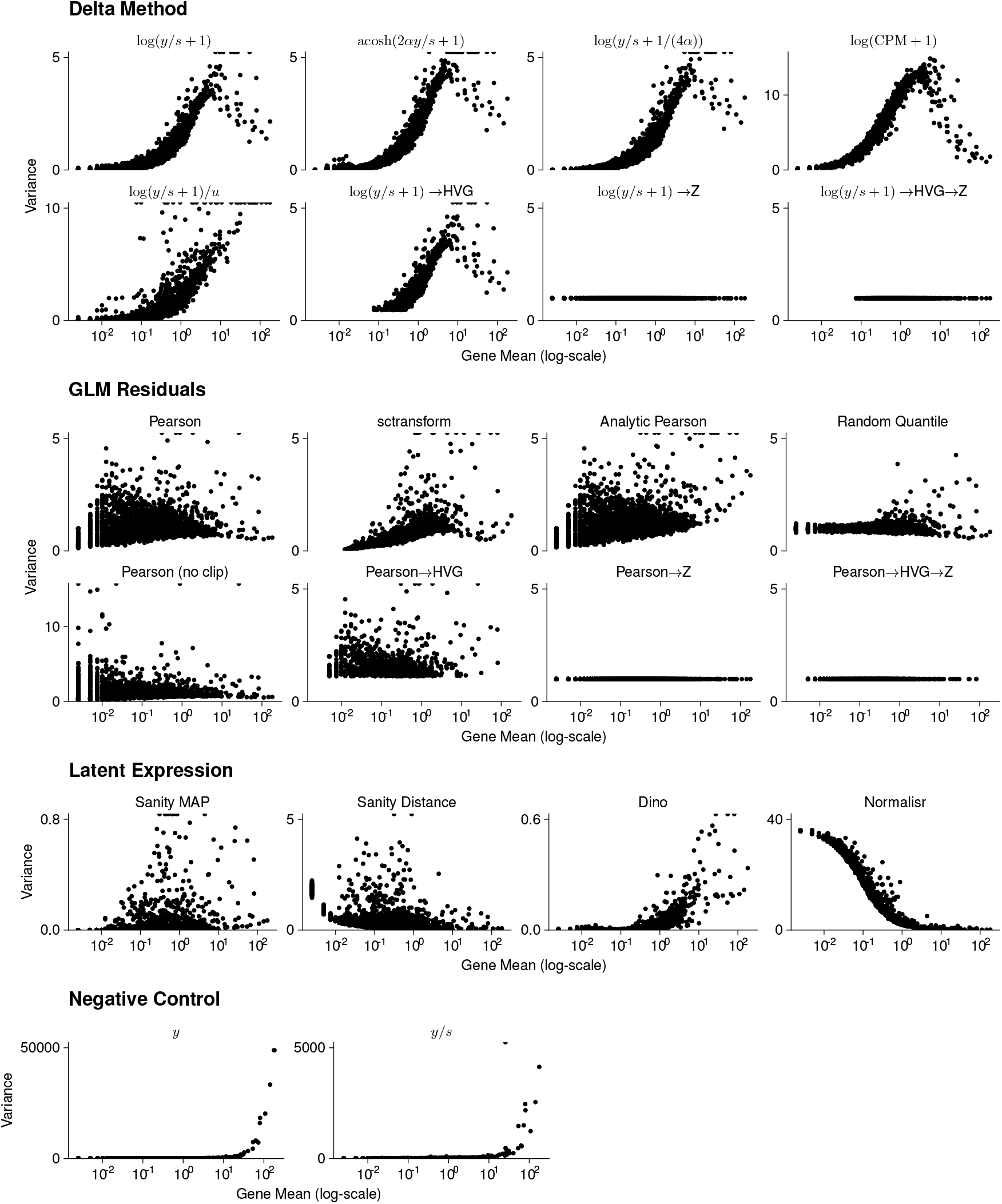
Scatter plot of the variance per gene after applying the variance-stabilizing transformation against the means of the *10X human hematopoietic cell* dataset subset to 400 cells and 5000 genes. Note that the scale of the y-axis differs for the raw counts, log(CPM + 1), log(*y/s* + 1)*/u*, Pearson (no clip), Sanity MAP, Dino, and Normalisr for aesthetic purposes. Points that exceed the y-axis scale are drawn on the top of each facet.

**Suppl. Figure S3:**
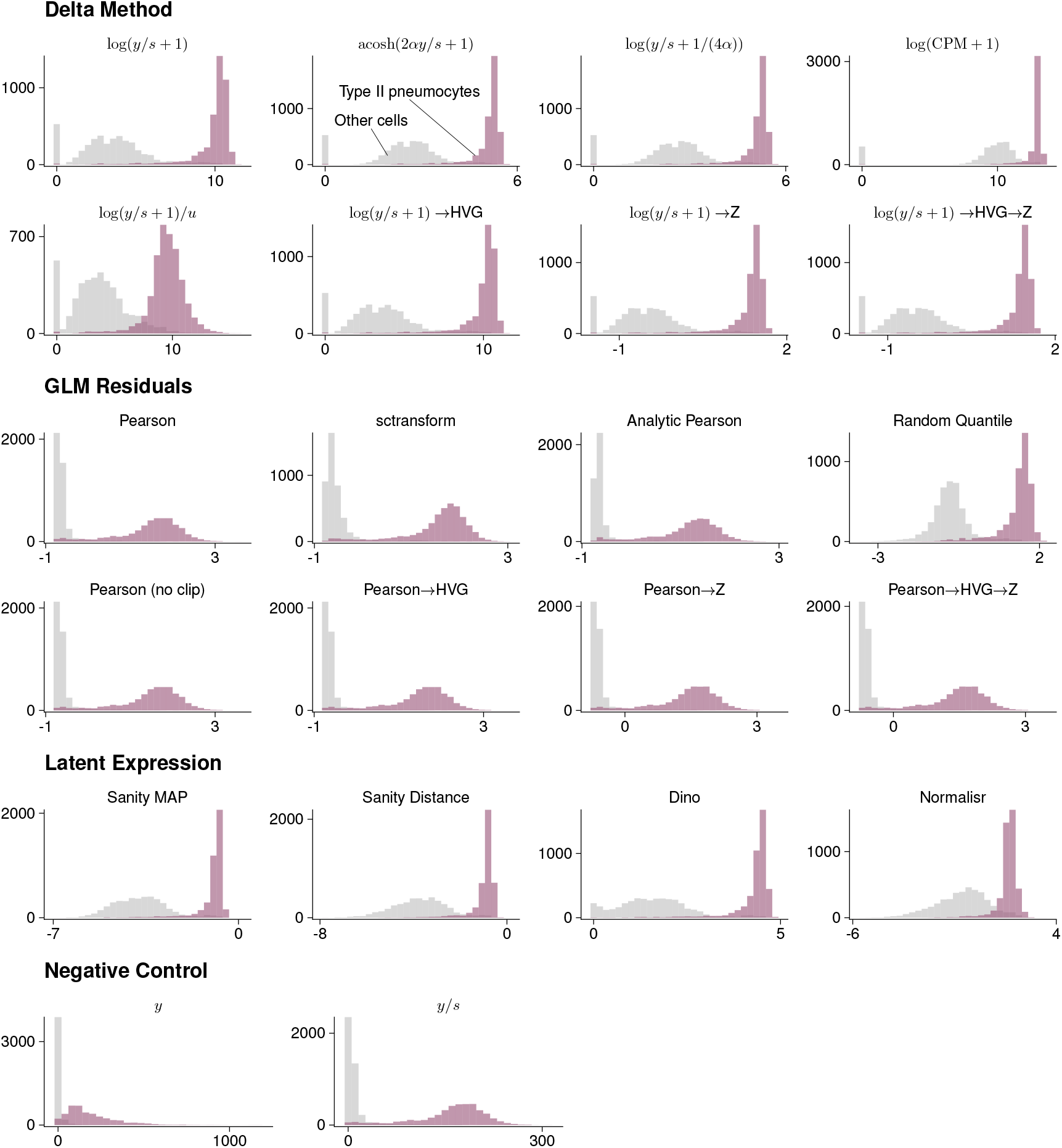
Histograms of the transformed values for a gene with a bimodal expression pattern. Counts from cells identified as type II pneumocytes are shown in purple and a matching number of counts from all other cell types are shown in grey.

**Suppl. Figure S4:**
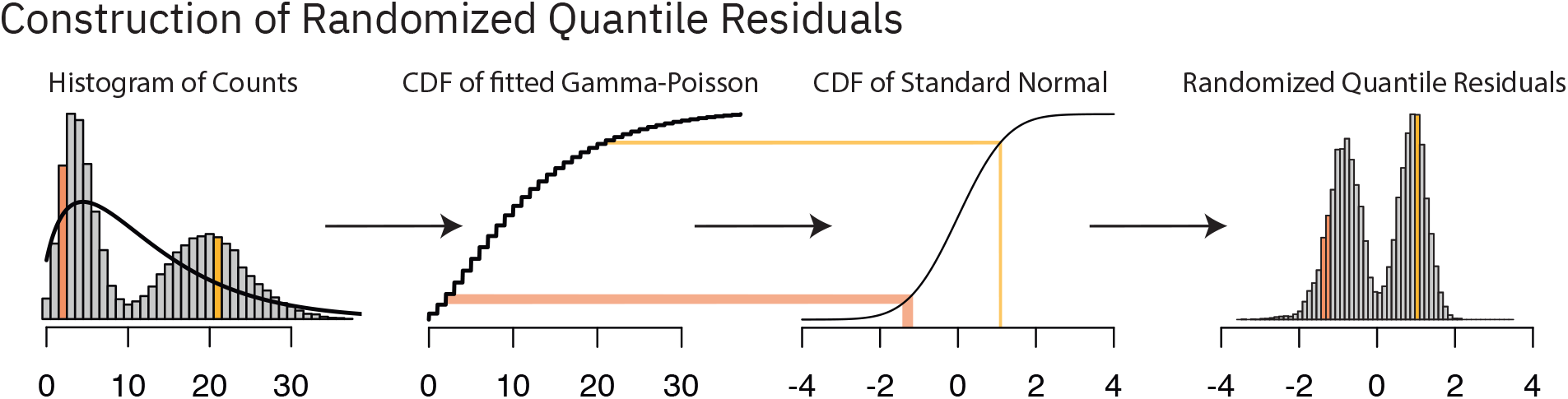
Schematic representation of how randomized quantile residuals are constructed. In the first step, a Gamma-Poisson distribution (black line) is fitted to the observed counts. Then, the quantiles of the Gamma-Poisson distribution are matched with the quantiles of a standard normal distribution by comparing their respective cumulative density functions (CDFs). This obtains a mapping from the raw count scale to a new, continuous scale. The two colored bars (orange for *y* = 2, yellow for *y* = 21) exemplify this mapping. The non-linear nature of the CDFs ensures that small counts are mapped to a broader range than large counts. This helps to stabilize the variance on the residual scale. Furthermore, the randomization within the mapping sidesteps the discrete nature of the counts.

**Suppl. Figure S5:**
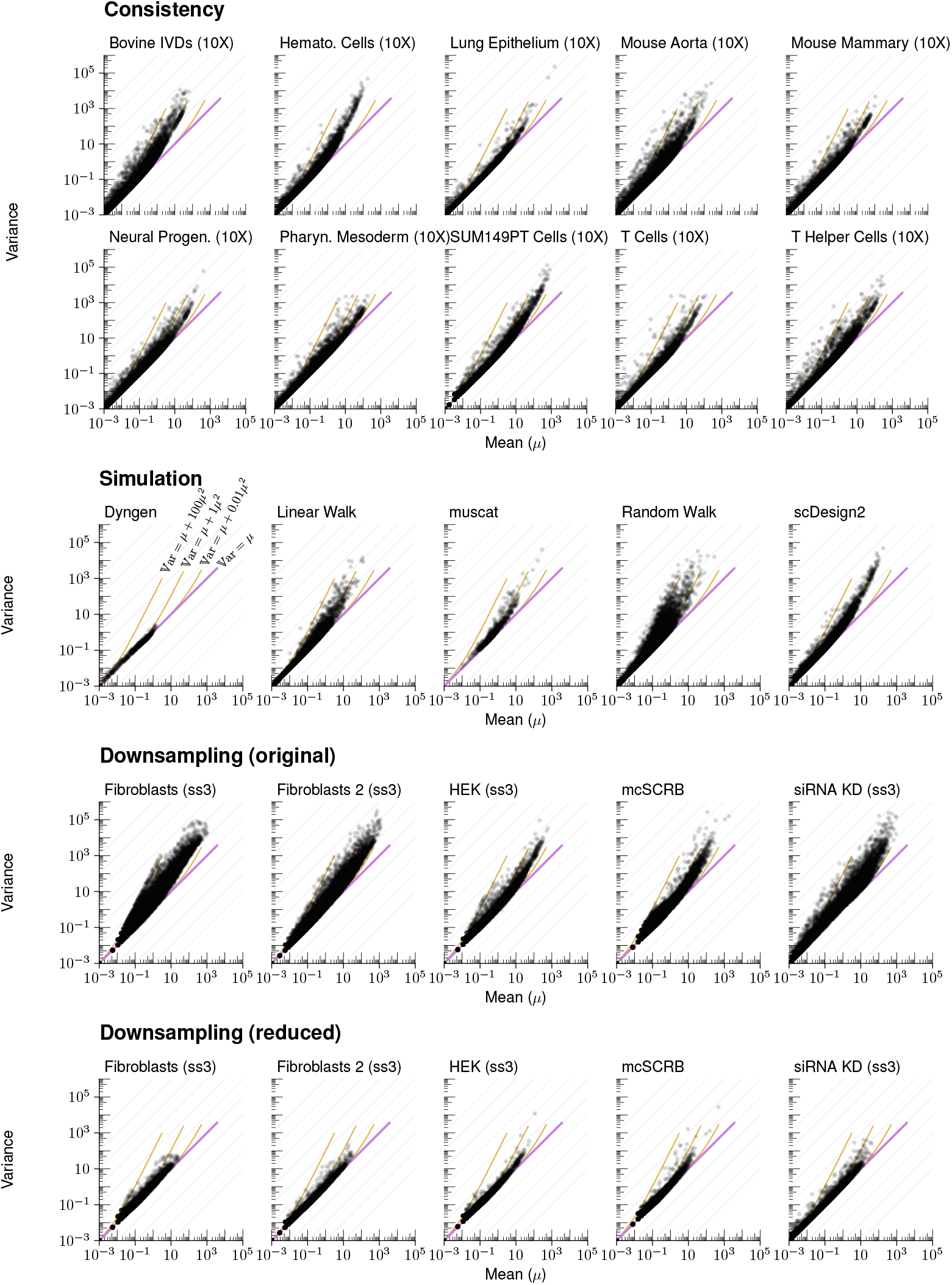
Log-log scatter plot of the mean-variance relation across all genes for each dataset. As size factor variations between cells introduce heterogeneity, for each dataset, we filtered out the largest and smallest 25% of cells.

**Suppl. Figure S6:**
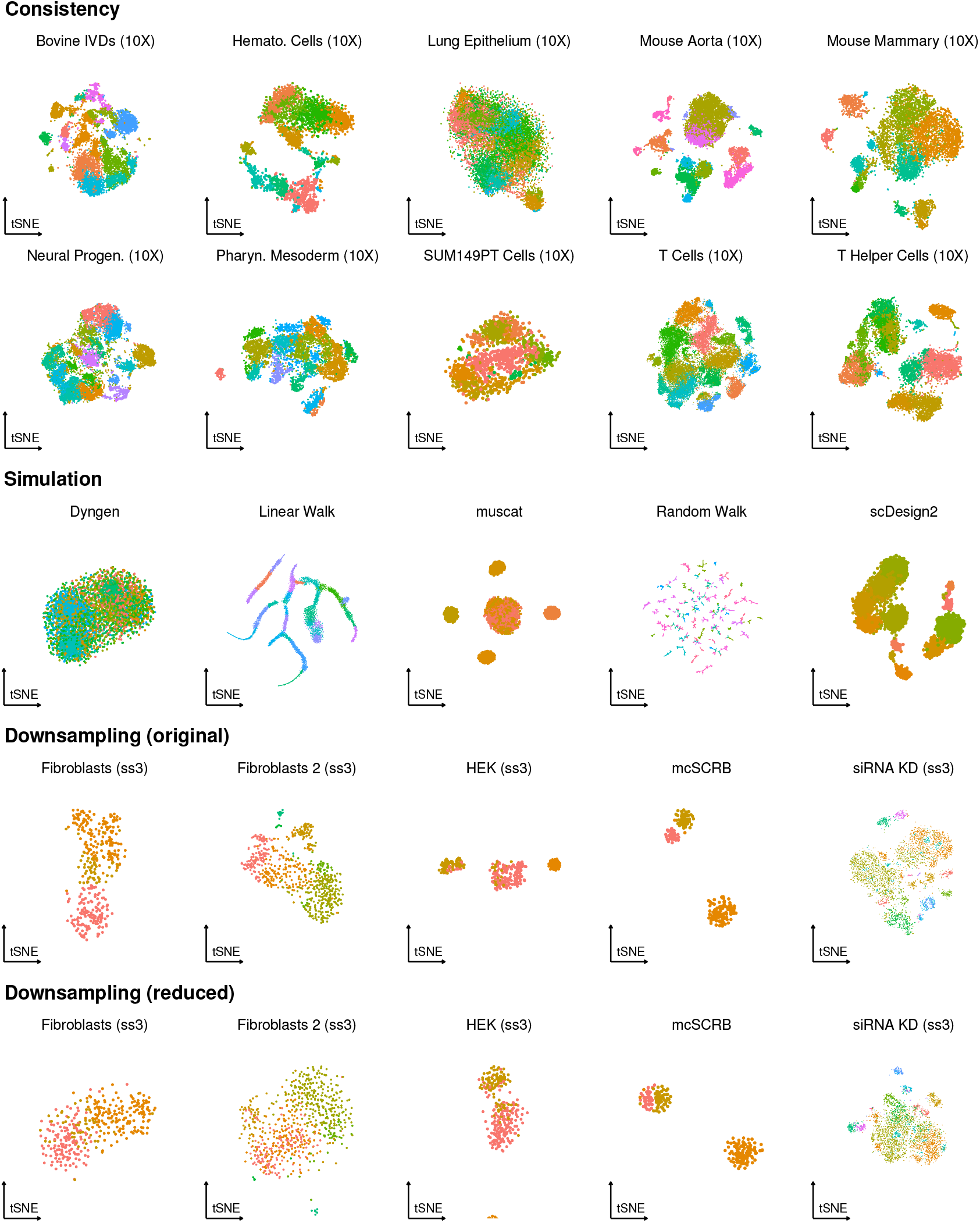
tSNE plot of each dataset. The cells are colored by clustering using the walktrap clustering algorithm. For the consistency data we clustered the counts after transformation with the shifted logarithm. For the simulation data, we clustered the ground truth. For the downsampling data, we clustered the deeply sequenced data after transformation with the shifted logarithm.

**Suppl. Figure S7:**
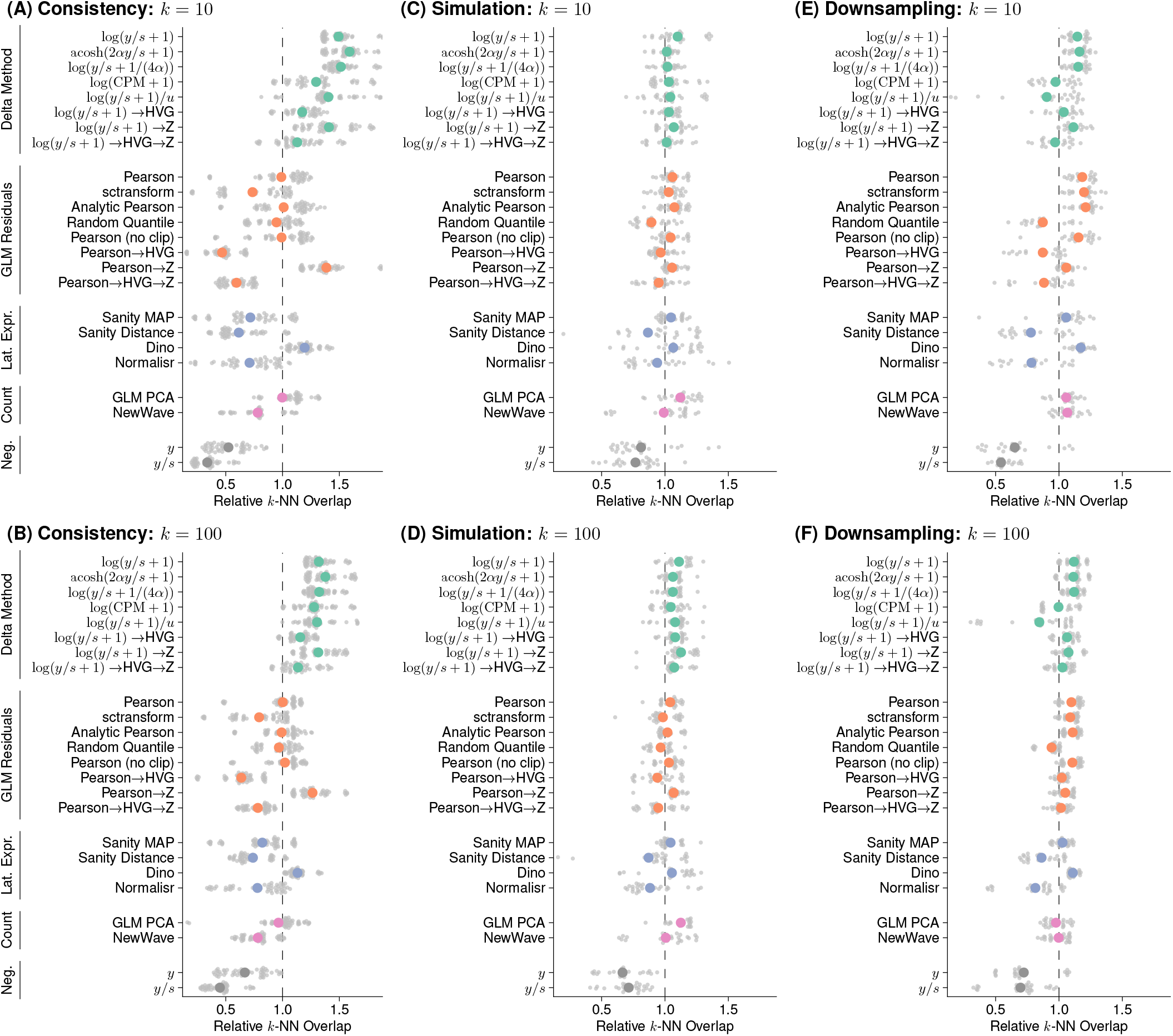
Plot of the aggregate results of the consistency (A, B), simulation (C, D), and downsampling benchmarks (E, F) for *k* = 10 and *k* = 100, respectively. The results for each dataset are broad to a common scale by normalizing to the mean *k* nearest neighbor overlap per dataset. The colored points show the mean per group.

**Suppl. Figure S8:**
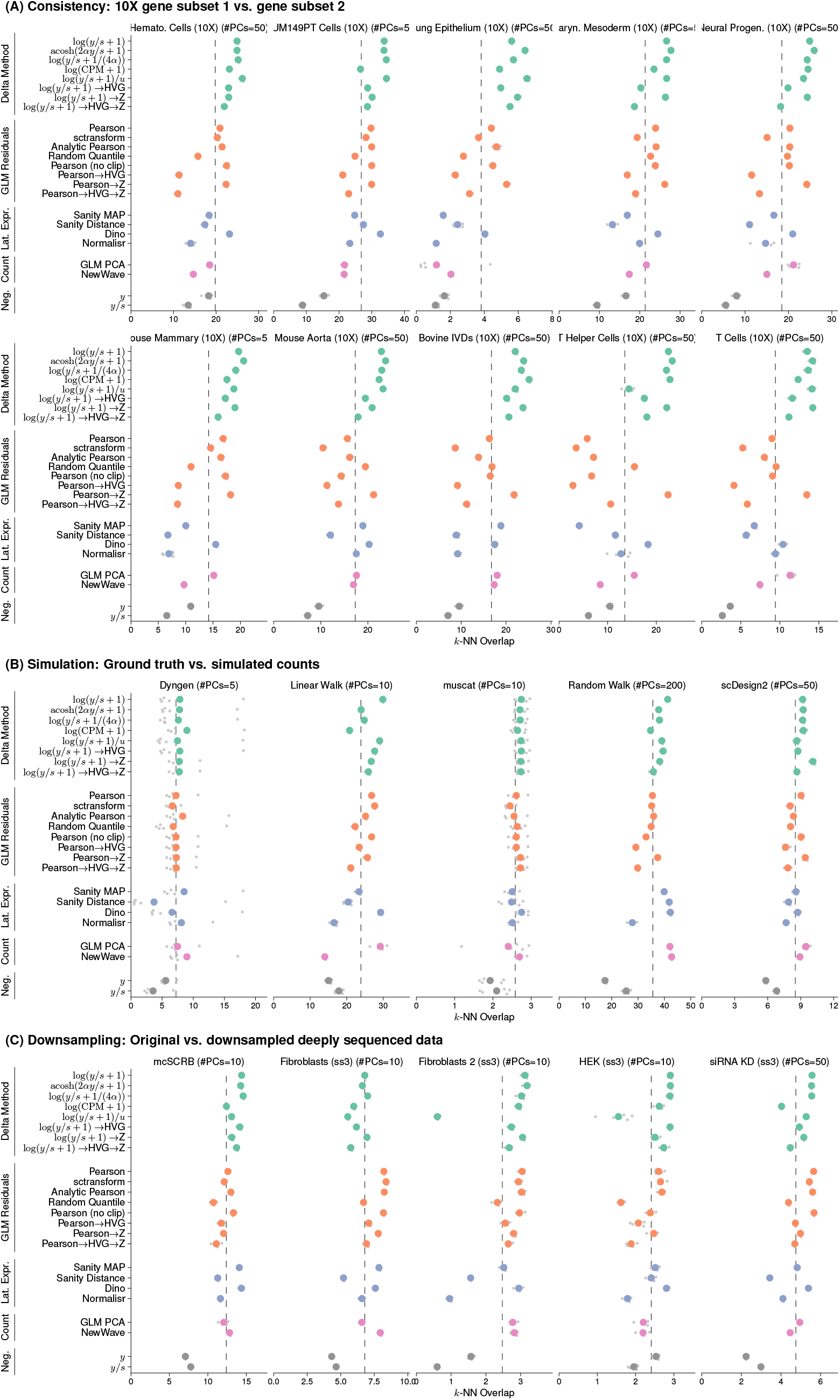
The unaggregated results from the consistency (A), simulation (B), and downsampling benchmarks (C) for *k* = 50. The grey points show the raw results from the five replicates per dataset; the colored points show their mean. The dashed vertical line indicates the mean *k*-NN overlap per dataset and is the reference used to aggregated the results as shown in Fig. 2A-C. The subtitles of each plot indicate the number of dimensions used for the PCA per dataset, which we chose based on the complexity of the dataset.

**Suppl. Figure S9:**
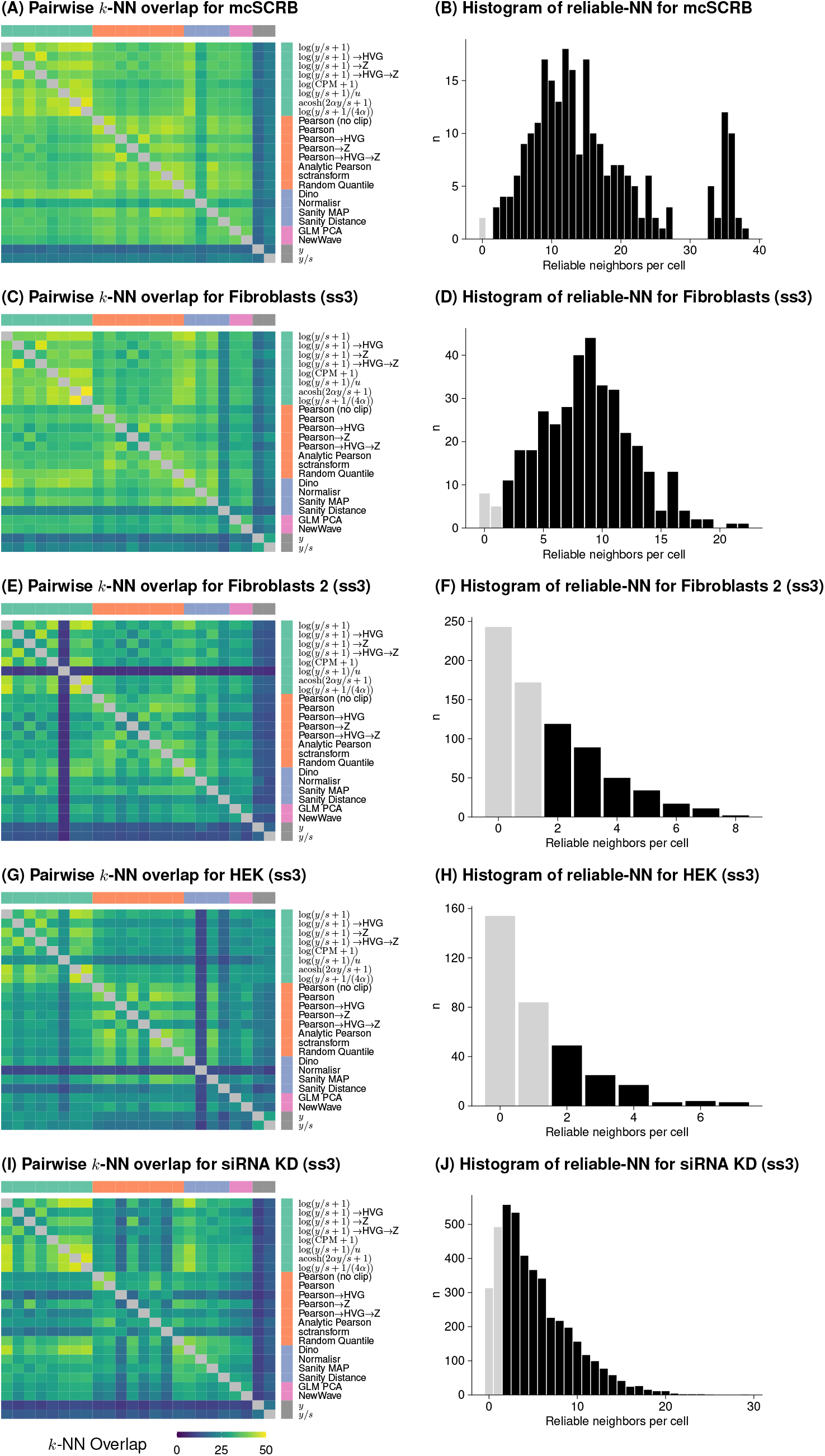
Inference of the reliable nearest neighbors for the five deeply sequenced datasets. The left column (A,C,E,G,I) shows heatmaps of the average *k*-NN overlap for all pairs of transformations. The right column (B,D,F,H,J) shows histograms of the number of reliable neighbors per cell (i.e., the neighbors among the 50 *k*-NN that were identified by all 22 transformations). The dark shaded bars show the cells that were used to calculate the overlap with the downsampled version of the data in Suppl. Fig. S8C.

**Suppl. Figure S10:**
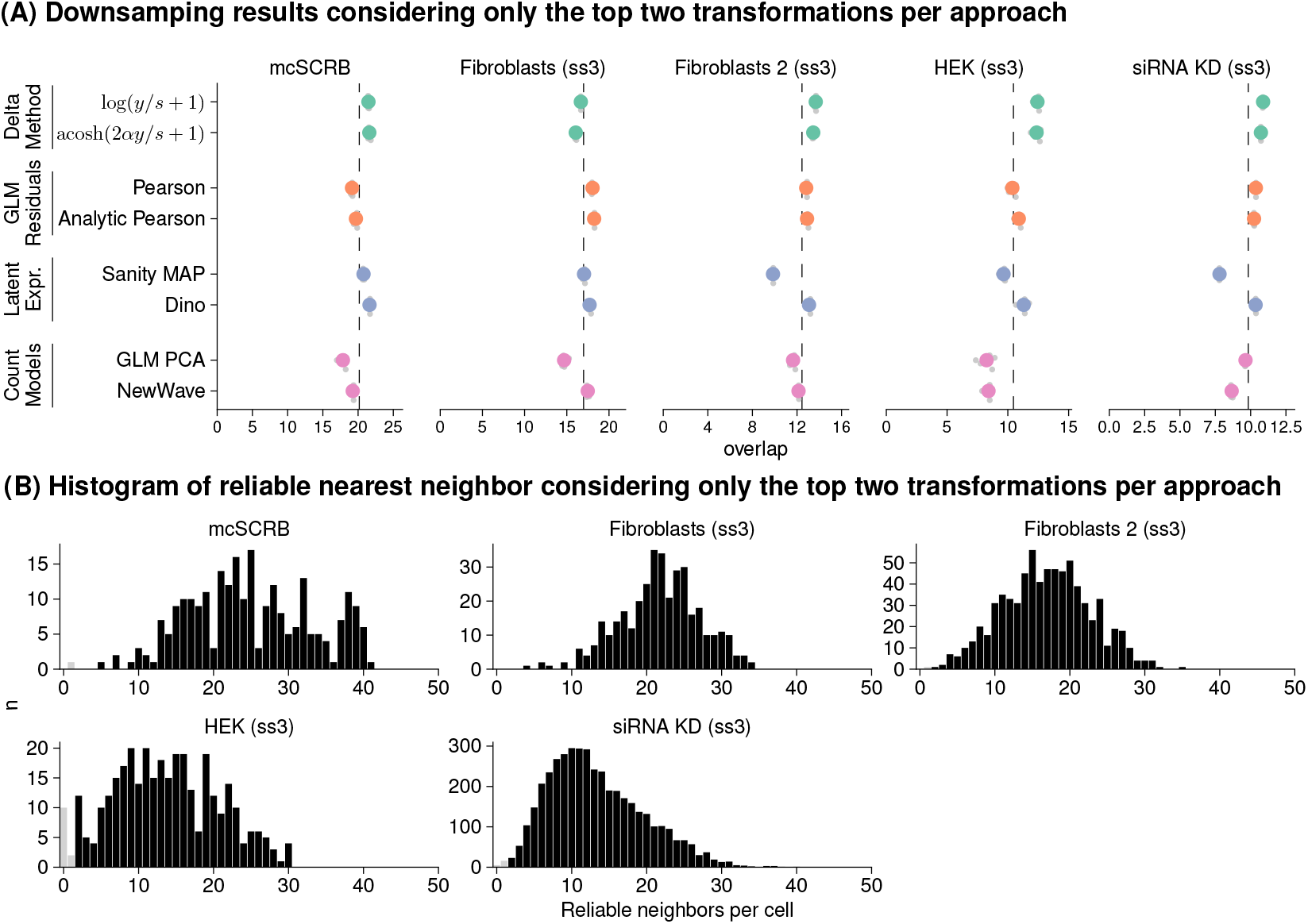
The downsampling benchmark considering only the top two transformations per approach. (A) The unaggregated results of for downsampling benchmarking using the same settings as in Suppl. Fig. S8C. (B) The histograms of reliable neighbors per cell only considering the eight transformations from (A).

**Suppl. Figure S11:**
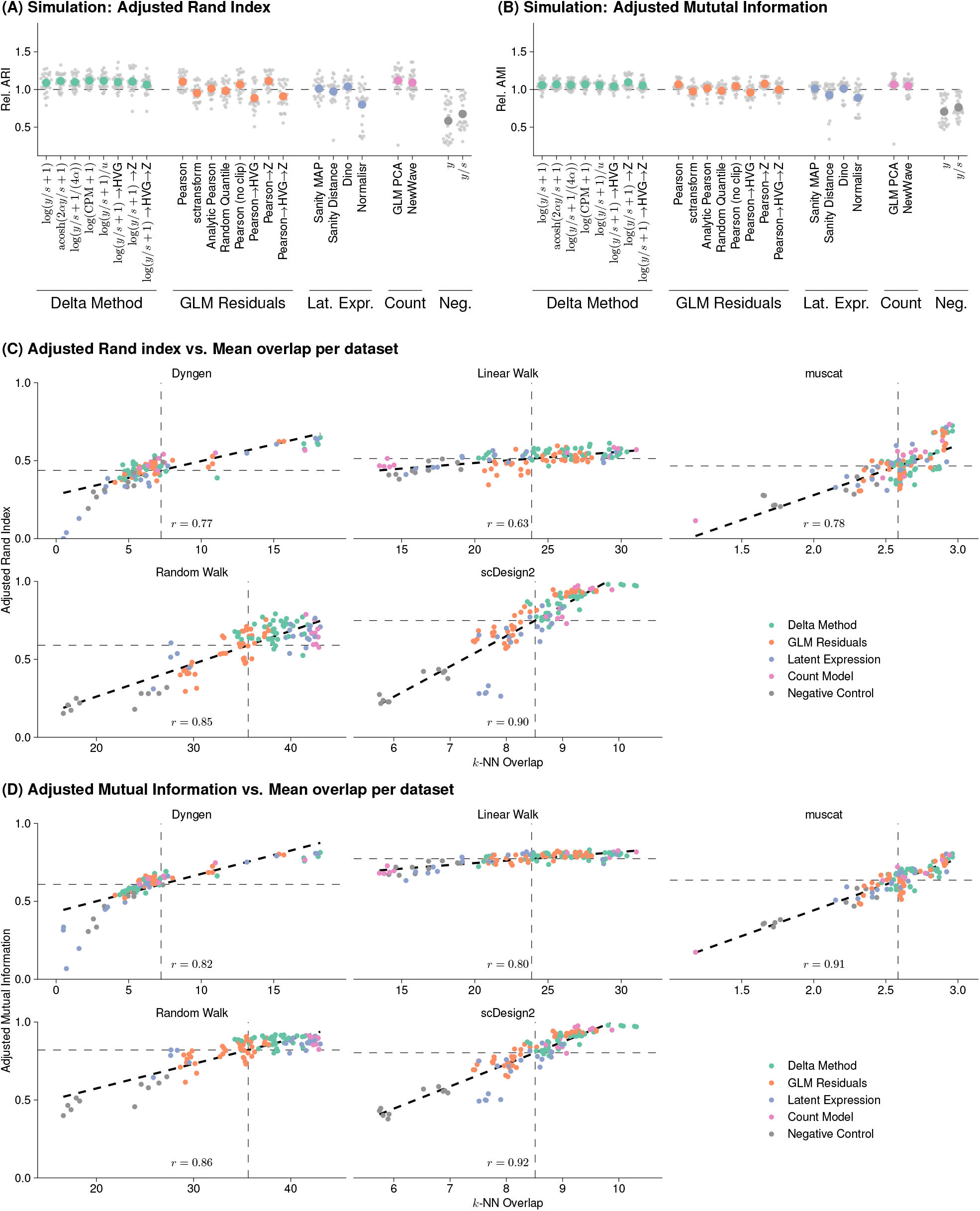
Results of the simulation benchmark in terms of the adjusted Rand index (A) and the adjusted mutual information (B) instead of the *k*-NN overlap. (C-D) Scatter plots facetted by simulation framework that compares the results for the *k*-NN overlap with the adjusted Rand index and adjusted mutual information, respectively. Each point is one replicate for the transformation results of that dataset colored by the transformation approach. The black dashed line shows the linear fit and the number at the bottom of each plot is the correlation coefficient. The horizontal dashed line is the mean ARI / AMI that is used for forming the relative performance in (A) and (B). The vertical dashed line is the mean *k*-NN overlap and corresponds to the vertical dashed line in Suppl. Fig. S8B.

**Suppl. Figure S12:**
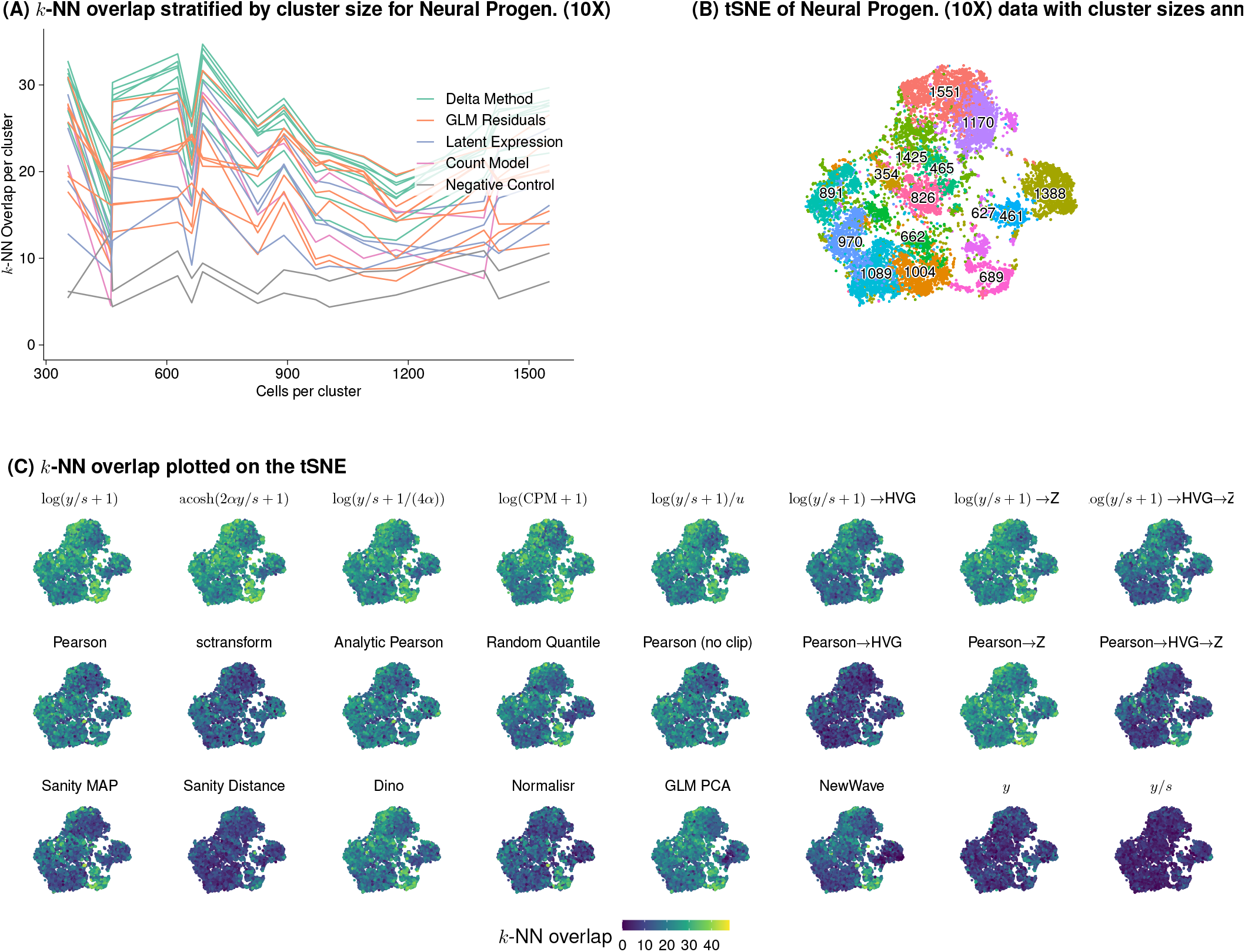
*k*-NN overlap of the two halves of the *human neural progenitor* dataset stratified by cluster (A, B) and by location in the two-dimensional tSNE projection (C).

**Suppl. Figure S13:**
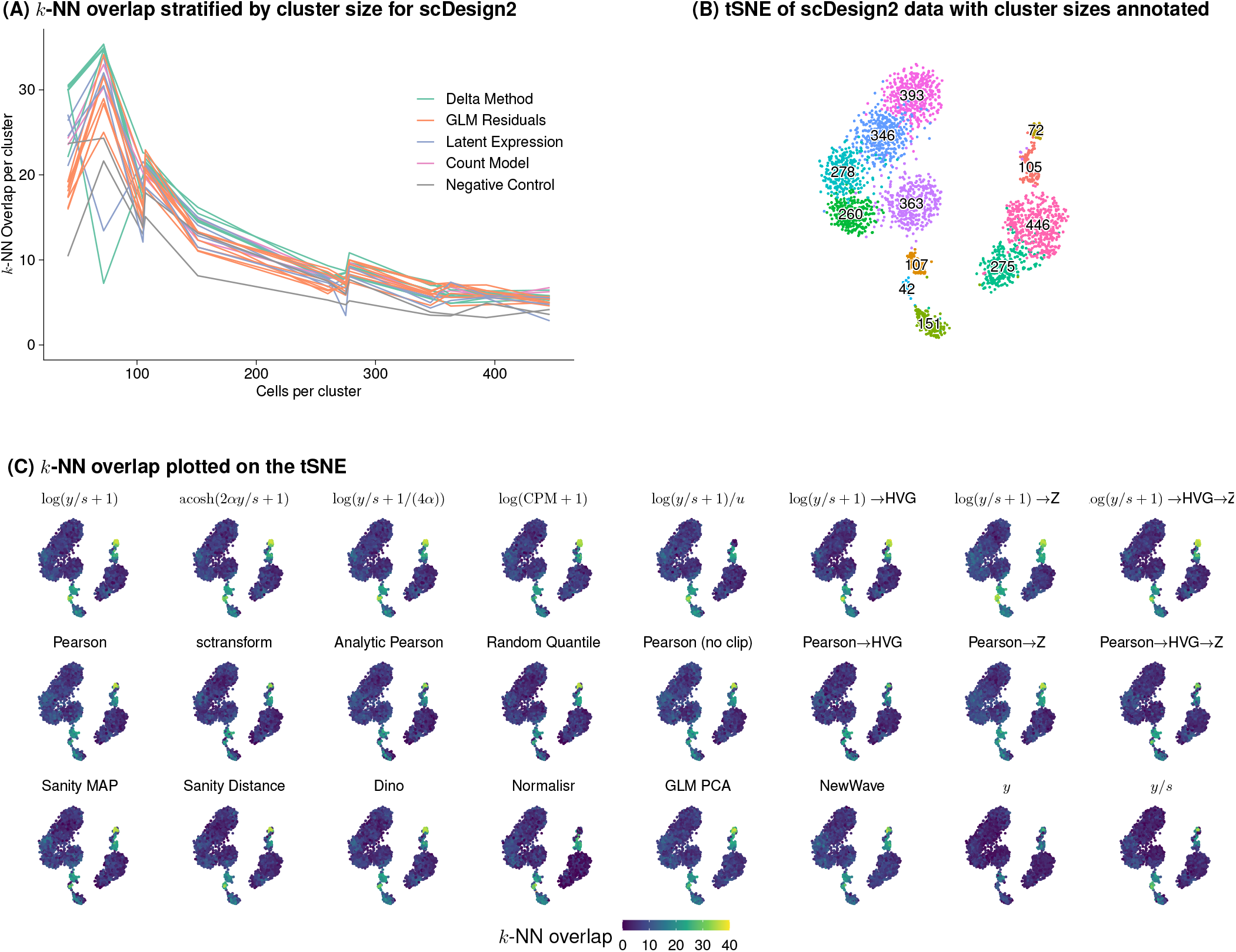
*k*-NN overlap on the dataset simulated with scDesign2 stratified by cluster (A, B) and by location in the two-dimensional tSNE projection (C).

**Suppl. Figure S14:**
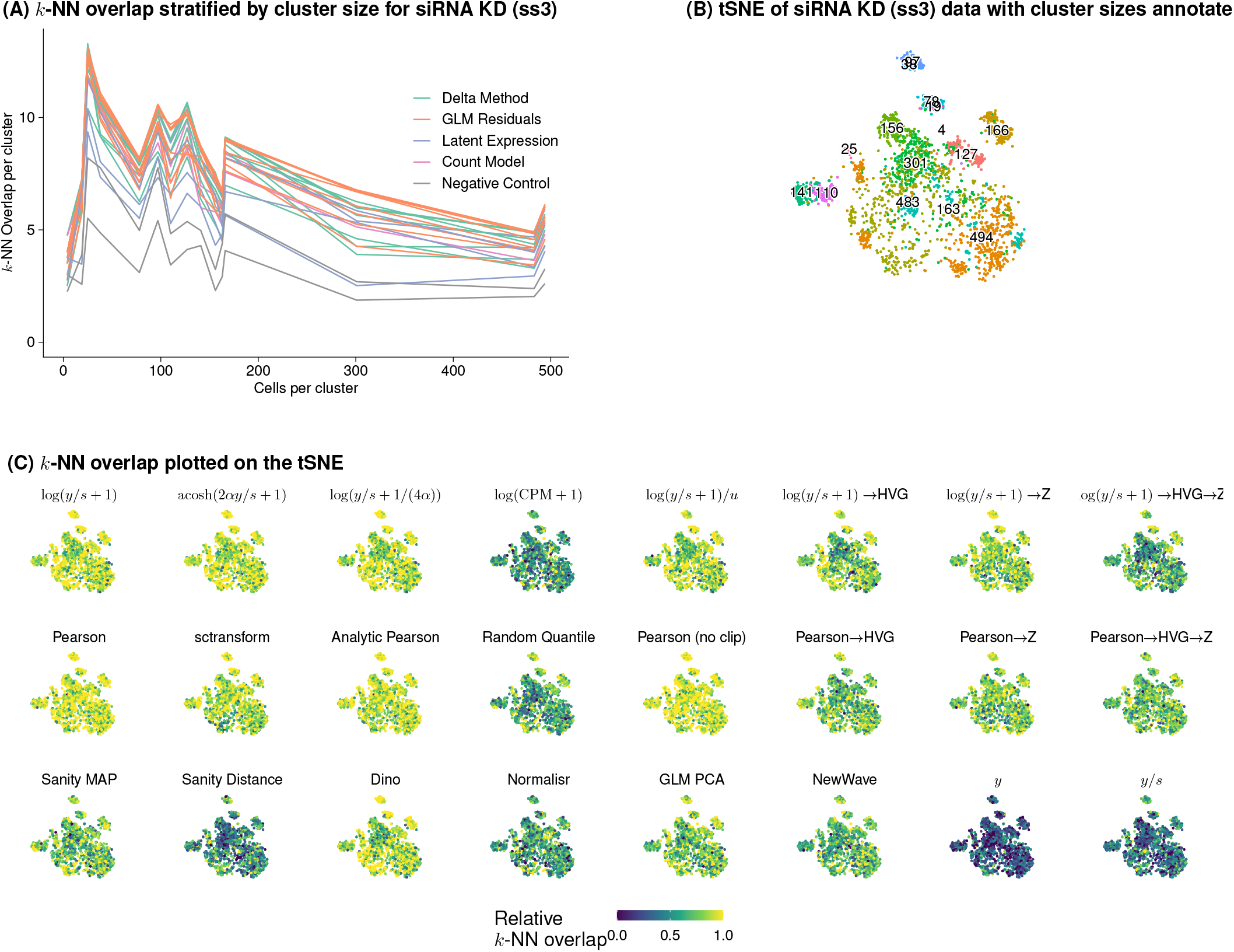
(Relative) *k*-NN overlap between the deeply sequenced and downsampled version of the siRNA knockdown dataset stratified by cluster (A, B) and by location in the two-dimensional tSNE projection (C). All cells for which the intersection of nearest neighbors on the deeply sequenced data was less than 4, were filtered out.

### B Appendix

#### B.1 Variance-stabilizing transformation for a quadratic mean-variance relation

The Gamma-Poisson distribution with mean *μ* and overdispersion *α* implies a quadratic mean-variance relation

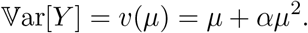

Our goal is to find a function *g* for which

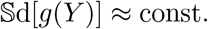

The delta method approximates the standard deviation of a transformed random variable as

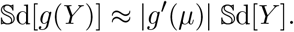

We can require this to be constant and solve for |*g*′(*μ*)|:

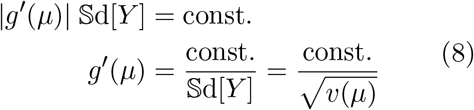

Given the derivative *g*′, we can use integration to identify the functional form of our transformation (note that without loss of generality, we can ignore the constant, whose value does not affect the variance stabilization property.)

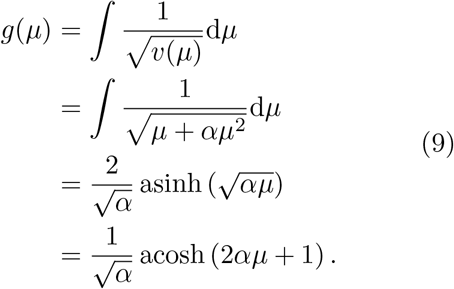

The last two expressions are mathematically equivalent. In the paper, we preferentially use the acosh-based expression since it seems slightly simpler. It is, however, worth noting that in the past, the name asinh transformation has been used (Bartlett, 1947).

If there is no overdispersion (*α* = 0), the acosh transformation reduces to the well-known square root variance stabilizing transformation for Poisson random variables

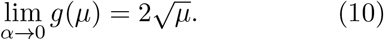

#### B.2 Approximating the acosh transformation with the shifted logarithm

The inverse hyperbolic cosine (acosh) transformation from Eq. (1) can also be expressed in terms of the logarithm function,

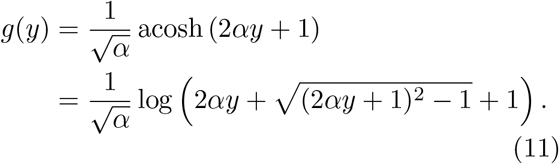

We want to approximate this transformation using the shifted logarithm and thus find *a, b*, and *c* in

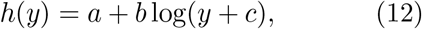

so that *h*(*y*) ≈ *g*(*y*).

We aim to find *a, b*, and *c* such that for large *y, h*(*y*) converges to *g*(*y*). We notice that

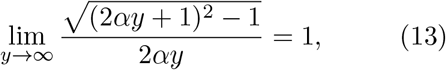

and thus for large *y*

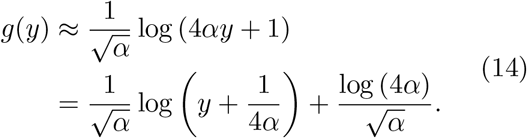

The linear scaling *b* and the offset *a* do not influence the variance stabilization; the important insight is that the pseudo-count 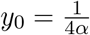 ensures that the shifted logarithm is most similar to the variance-stabilizing transformation derived using the delta method.

#### B.3 Delta method-based variance-stabilizing transformation and size factors

Suppl. Fig. S1 demonstrates that delta method-based variance-stabilizing transformations struggle to account for varying size factors.

To incorporate cell-specific size factors in the delta method-based variance stabilizing transformation approach, the counts *Y*_*ij*_ are divided by the size factor *s*_*j*_ before applying the transformation: *g*(*Y*_*ij*_*/s*_*j*_) (Love et al., 2014). To see the implications of this, it is helpful to look at a decomposition of the variance of a Gamma-Poisson random variable *Y* :

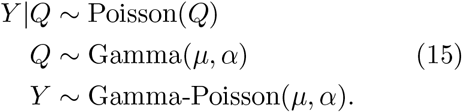

In the context of RNA-seq count data, the Poisson level of this hierarchical model represents the technical sampling noise and *Q* models additional variation. According to the law of total variation

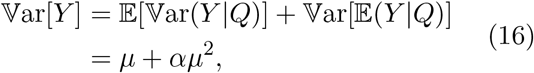

where 𝕍ar[*Y* |*Q*] = *μ* and 𝕍ar[*Q*] = *αμ*^2^.

If we apply the same approach to a model with size factors

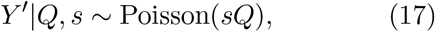

we find that

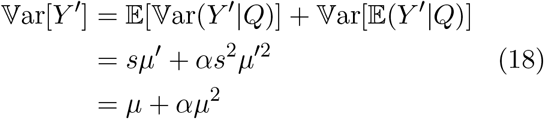

where *μ* = *sμ*′.

If, however, we want to apply the delta method-based variance-stabilizing transformation to a size factor standardized count

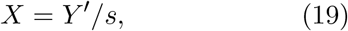

we find that

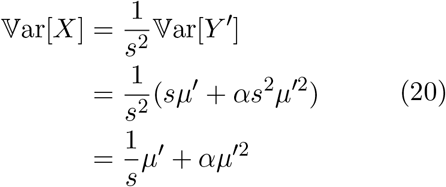

The difference between the final line of Eq. (18) and Eq. (20) explains the problem observed when applying the delta method-based variance-stabilizing transformation to correct data where the size factors vary a lot between cells.

## Footnotes

1 also referred to as the Negative Binomial distribution

